# Human knee osteoarthritis patient-specific cartilage-on-a-chip model captures donor differences to stressors and treatments

**DOI:** 10.64898/2026.09.03.748852

**Authors:** Lauren Banh, Kevin Perera, Brendan R. Lobo, Kara Walz, Ka Kit Cheung, Sutirtha Chatterjee, Esme Bonnell, Kebin Li, Byeong-Ui Moon, Rajiv Gandhi, Edmond W. K. Young, Sowmya Viswanathan

**Affiliations:** Osteoarthritis Research Program, Division of Orthopedic Surgery, Schroeder Arthritis Institute, University Health Network, Canada; Krembil Research Institute, University Health Network, Canada; Institute of Biomedical Engineering, University of Toronto, Canada; Department of Mechanical & Industrial Engineering, University of Toronto, Canada; National Research Council Canada, Boucherville, QC, Canada; Department of Materials Science & Engineering, University of Toronto, Canada; Division of Hematology, Department of Medicine, University of Toronto, Canada

## Abstract

Knee Osteoarthritis (KOA) is a progressive whole-joint disease without approved disease modifying OA drugs (DMOADs). Effective treatments have been hindered by multiple layers of heterogeneity, including diverse disease etiology and patient-to-patient variability. Here, we present a scalable, KOA patient-derived (PD) cartilage-on-a-chip (CartChip) model integrating end-stage KOA cartilage tissue explants on a microengineered platform that, under mechanical overloading and hyperinflammatory stressors, mimics different KOA etiologies. These stressors drove distinct multivariable model features, including changes in a curated panel of anabolic and catabolic genes, extracellular matrix protein and soluble factors. Exploratory analysis of coordinated model readouts identified stressor-agnostic and -specific KOA disease signatures. Despite using KOA tissue, the model showed improvements to dexamethasone, a symptom-modifying, anti-inflammatory KOA treatment. The model responses to dexamethasone were dependent on both stressor and donor heterogeneity. Exploratory groupings of coordinated model readouts provided proof-of-concept for predicting categories of patient responsiveness to test therapeutics. Annotating patient data provided additional donor-dependent contexts for interpreting model responsiveness. PD-CartChip provides a powerful research platform to potentially surmount the donor and stressor heterogeneity barrier in developing DMOADs.

## Introduction

Osteoarthritis (OA) is a degenerative whole joint disease affecting over 600 million people globally and is a leading cause of disability worldwide^1^. Despite its high prevalence, current therapeutic interventions are unable to effectively halt long-term joint destruction. Treatment strategies for OA are highly stage-dependent and as the disease progresses from minor degeneration to total joint destruction, the treatment approach shifts from prevention and lifestyle management to invasive surgical interventions including total knee arthroplasty (TKA)^2^. The limited success of current therapies stems partly from many disease-modifying OA drug (DMOAD) trials adopting a “one-size-fits-all” approach for a highly complex, heterogeneous disease^3,4^. Patient heterogeneity, however, is difficult to capture, and achieving effective patient stratification in preclinical therapeutic testing models and clinical trial designs remains a challenge, despite its known importance to disease progression and treatment responses.

It is well established that patient heterogeneity reflects a continuous biological spectrum^5^. Within this landscape, phenotypic and molecular endotypic classification schemes^6,7^ have emerged and demonstrate how different etiological origins of knee OA (KOA) converge into overlapping clinical presentations. While animal models provide necessary dynamic insights^8^, they cannot recapitulate human-specific disease complexity in an easily scalable manner.

Advanced microfluidic bioreactors known as “cartilage-on-chips” (COC) have emerged as promising models for combining microscale engineering with biological relevance. Recent COC platforms^9–14^ have demonstrated that mechanical actuation can be integrated with microfluidics and 3D cell culture to capture mechanical dysfunction, a major driver of cartilage degradation, through load-based mechanisms. However, no single model can collectively recapitulate distinct pathogenic drivers, disease-specific stages and responses, and inter-patient variability^15,16^. Together, these considerations highlight an unmet need for COC platforms that combine microscale precision with structural and biomechanical relevance to native cartilage with multiplexing capabilities. By prioritizing the use of native patient-specific cartilage testing, such systems could more faithfully reproduce the mechanobiological drivers of OA and improve the predictive power gap left by existing *in vitro* models.

Here, we describe our novel “PD-CartChip” system, which is the first COC model to incorporate human KOA patient-derived cartilage explants, using a flexure mechanism capable of delivering dynamic cyclic compressive forces across a parallel array of explant samples with varying dimensions. Using our system, we demonstrate differences between mechanical and pro-inflammatory stressors in driving cartilage degradation and matrix remodelling and show an ability to dually capture patient-specific and stressor-dependent responses to dexamethasone (DEX), representing a standard-of-care therapeutic class used to manage KOA symptoms.

## Results

### PD-CartChip and experimental workflow

PD-CartChip allows, for the first time, controlled physiological mechanical loading on human cartilage biopsies, enabling functional interrogation of millimetre-scale patient-derived explants. The PD-CartChip system (Fig. 1) consists of: (i) a linear actuator mounted on a base plate with associated brackets, (ii) a consumable plastic flex device with flexure mechanisms and an array of compartment wells for housing the cartilage explant samples, and (iii) the patient-derived cartilage explants themselves. The linear actuator and plastic flex device constitute our “FlexChip” platform, which is tissue-agnostic and capable of applying mechanical loading of various modes to any 3D tissue sample of appropriate size, including tissue-engineered hydrogel-based constructs. Once a patient-derived cartilage explant is loaded into our FlexChip platform, our PD-CartChip model, specific to cartilage, has been formed.

**Fig. 1.**
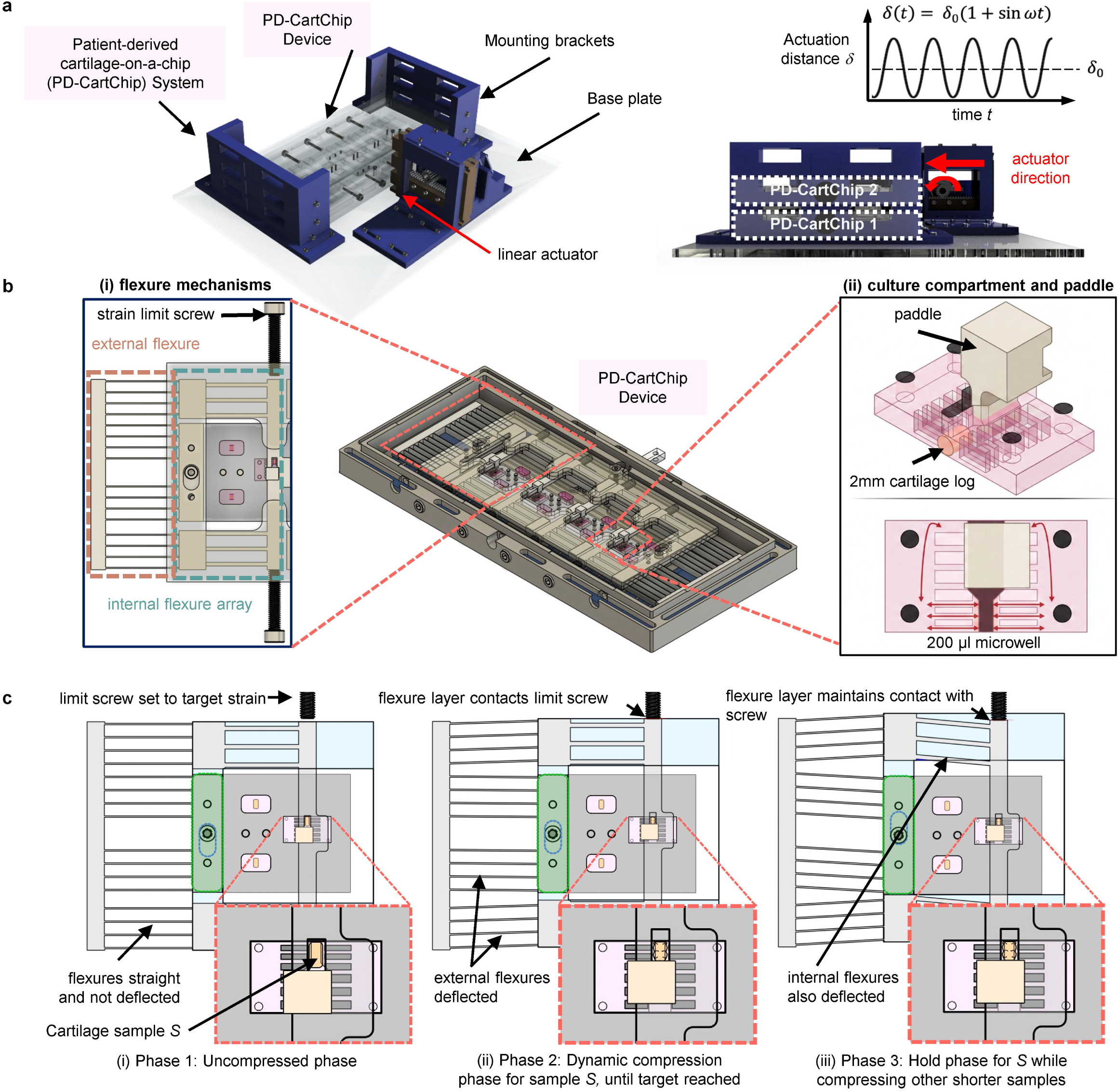
Overview of PD-CartChip system. **a,** The PD-CartChip system is comprised of the core PD-CartChip device and the actuation platform comprised of a base plate, mounting brackets, and a rack-and-pinion linear actuator (i.e., the tissue-agnostic FlexChip platform). One PD-CartChip system can currently accommodate two stacked PD-CartChip devices with four replicates per device. **b,** (i) Within a PD-CartChip device, an internal flexure array comprised of parallel acrylic beams carries the paddles to a targeted hard-stop strain limit screw for individual samples, as the actuator completes a full sinusoidal cycle. Symmetric external flexures ensure linear translation of all the paddles. (ii) ∼2 mm diameter cartilage logs are placed in the culture compartment and exposed to unconfined compression via the acrylic paddles. This microwell possesses 500-µm side channels and 0.3-mm clearance on either side of the cartilage logs to allow for liquid displacement during compression. **c,** During compression, the actuator drives a carriage frame where the internal flexure arrays are mounted. Each replicate goes through three phases: (i) an uncompressed phase where the flexures are straight and the cartilage is not under load; (ii) a dynamic compression phase, where the paddle and its flexure assembly compress an individual sample *S* to target strain (constrained by limit screw); (iii) a hold phase, where sample S maintains the target strain while other smaller samples continue Phase 2 to reach their specific strain target.

In this study, we applied cyclic compressive loading at physiologically relevant frequencies (1 Hz) and compressive strain magnitudes of up to 30% in PD-CartChip, while maintaining stable tissue positioning and reproducible deformation throughout extended culture periods. Parallelization of our system was enabled by compliant acrylic flexure beams within each device, allowing independent strain targeting of four samples per device. Stacking of multiple devices within our platform further increases the throughput (Fig. 1a,b). The device architecture is compatible with standard incubator environments and real-time stereomicroscopy to track explant compression, enabling long-term studies under controlled loading conditions. Furthermore, samples can be easily extracted from the device for downstream biomolecular analyses at the conclusion of the experiment. Paired with short- and long-term biological readouts, PD-CartChip offers a versatile longitudinal platform to understand disease progression alongside immediate and lasting treatment effects.

Leveraging this capability, we established an experimental workflow to streamline testing of freshly resected tissue from human KOA patients during TKA surgery (Fig. 2). Full-depth cartilage explants were loaded into PD-CartChip devices and either exposed to cyclic compressive loading to mimic mechanical compression acting on the knee joint or not exposed to any loading to model different static (uncompressed) test conditions.

**Fig. 2.**
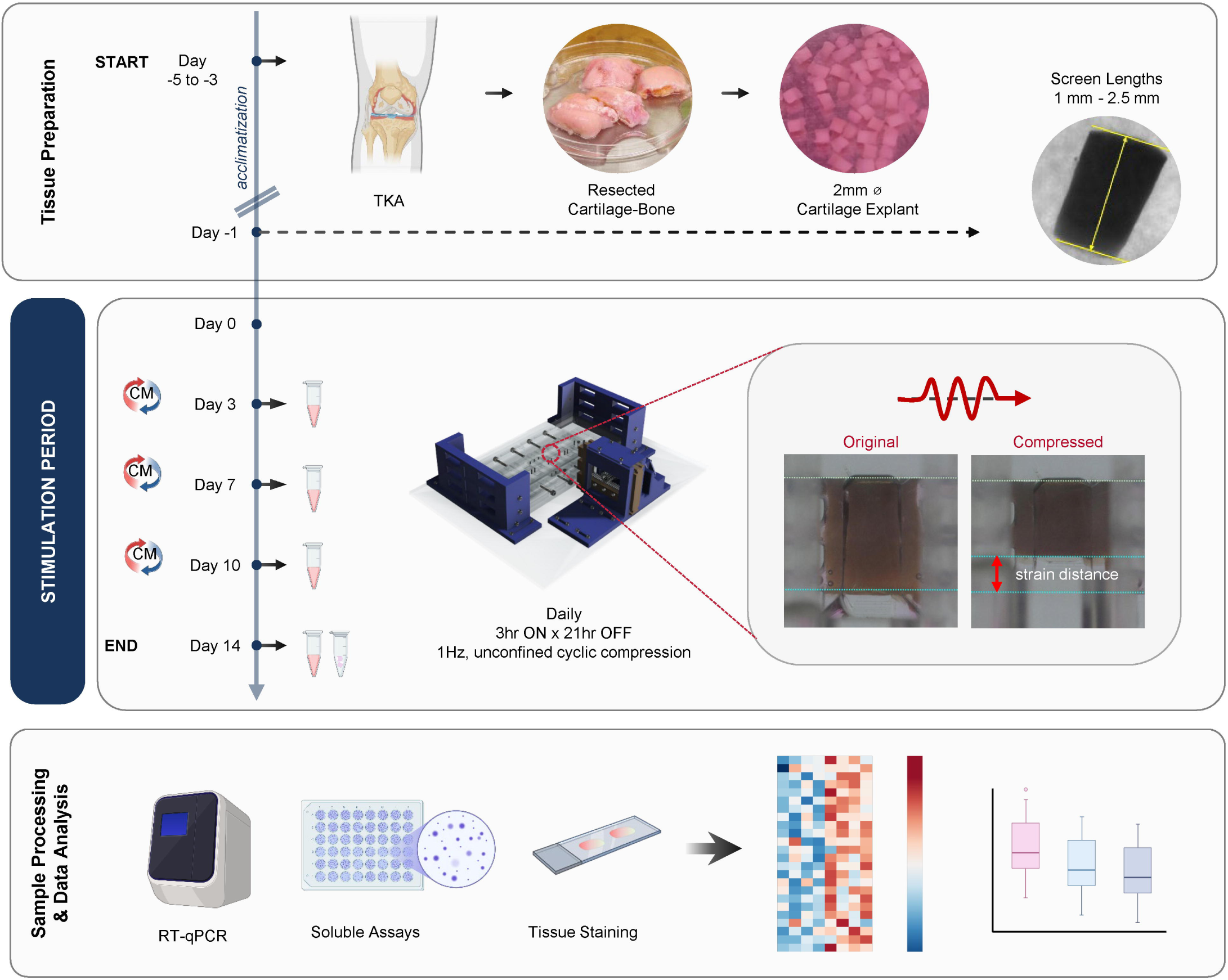
PD-CartChip model experimental pipeline. Human articular cartilage explants are acquired from total knee arthroplasty (TKA) surgery, processed into 2 mm diameter biopsy logs, acclimatized for 3-5 days, and screened by length (1 mm to 2.5 mm) prior to the experiment. Cartilage explants are then loaded into the device and either subjected to unconfined cyclic strain at a natural walking loading regime (1Hz, 3hr stimulation-21hr rest per day) or cultured in static culture conditions for 14 days. Supernatant (conditioned medium, CM) is collected and refreshed every 3-4 days before harvesting the supernatant and tissue at the end of the experiment. Post-experimental analysis includes real-time quantitative polymerase chain reaction (RT-qPCR), soluble factor assays, and tissue staining. Created with Biorender.com.

### Matrix degradation in cartilage is load-dependent

We first applied the PD-CartChip model and our experimental workflow to show how mechanical strain influences cartilage degradation in different patient-derived samples. To evaluate this, we compared patient-specific cartilage explants subjected to physiological compression (PC, 10% compressive strain) versus hyperphysiological compression (HPC, 30% compressive strain) against static unstimulated controls. Explants were compressed for 14 days using a standard regimen that mimics normal walking loads. Cartilage degradation was assessed by extracellular matrix (ECM) depletion, proteolysis-associated soluble factors, and gene expression profiles. We showed that sulfated glycosaminoglycan (sGAG) release was comparable across all conditions at day 7 (d7) but became significantly elevated with HPC compared to static conditions by day 14 (d14; Fig. 3a; Extended Data Fig. 1a). This was accompanied by tissue-specific Safranin-O staining that revealed an increasing trend in proteoglycan loss with HPC relative to PC and control (Fig. 3b,c; Extended Data Fig. 1b), suggesting potential low-level structural degradation.

**Fig. 3.**
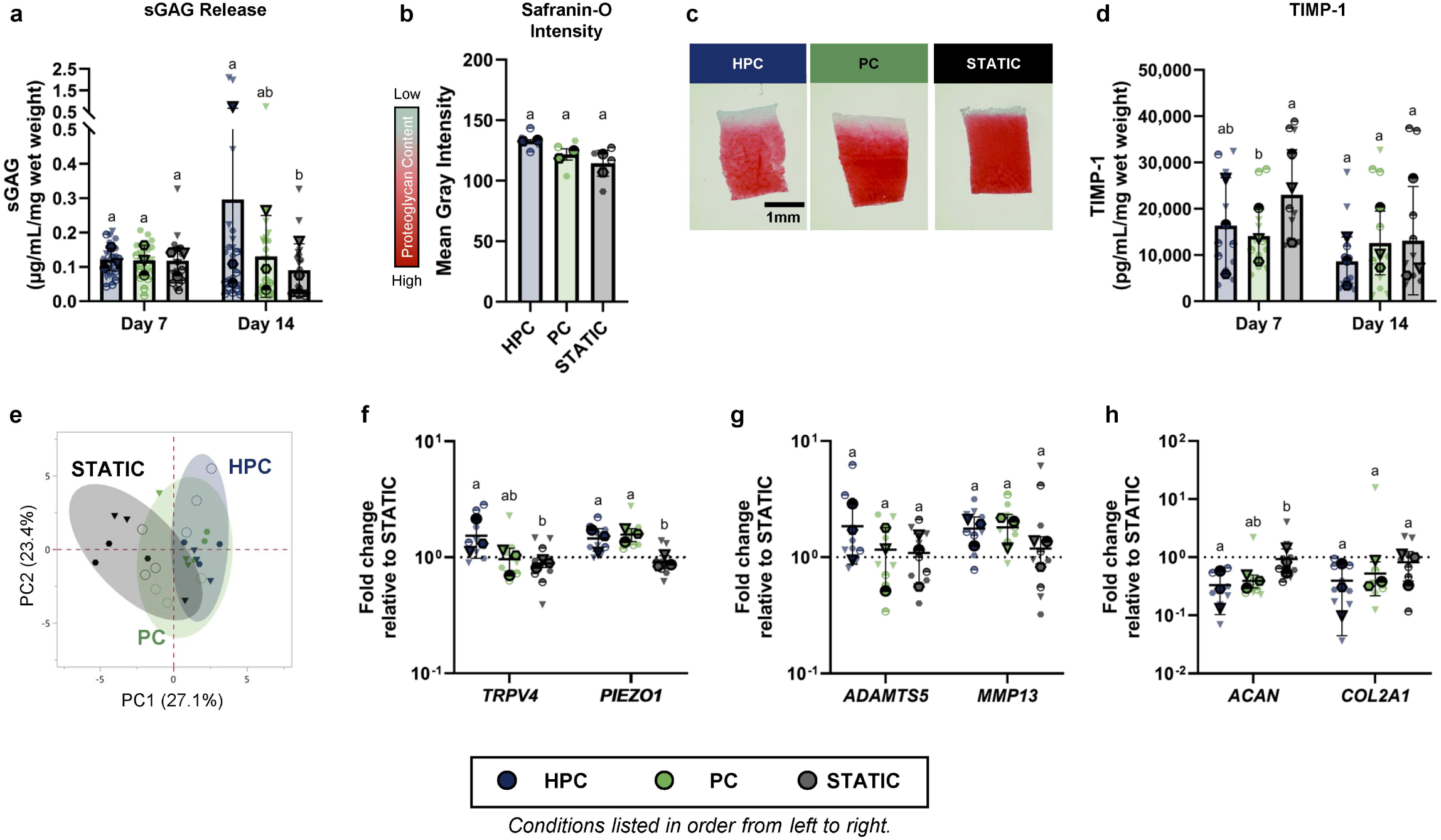
Hyperphysiological loading (HPC) induces differences in cartilage matrix level response compared to physiological loading (PC). **a,** Sulfated glycosaminoglycans (sGAG, µg/mL) release from the tissue to supernatant normalized to initial tissue wet weight (mg). **b,** Quantification of Safranin-O/Fast Green staining by mean gray intensity, where higher values indicate greater proteoglycan loss. **c,** Representative histological images of Safranin-O/Fast-Green stained cartilage sections at Day 14 where intense red staining corresponds to rich proteoglycan content, while a reduction in red intensity indicates matrix degradation. Scalebar = 1mm. **d,** Soluble tissue inhibitor of matrix metalloproteinases-1 (TIMP-1, pg/mL) levels normalized to initial tissue wet weight (mg). **e,** Principal component analysis (PCA) illustrating 2D clustering of individual samples based on donor-normalized gene expression (-ΔΔCT) across principal component 1 (PC1, 27.1% variance) and principal component 2 (PC2, 23.4% variance). Shaded regions represent 95% confidence ellipses for each respective group. Donor-blocked relative fold change in cartilage gene expression at Day 14 across **f,** mechanosensitive ion channels *(TRPV4, PIEZO1),* g, catabolic enzymes (*ADAMTS5, MMP13*), and **h,** cartilage-specific *(ACAN, COL2A1)* markers, calculated via 2^-ΔΔCT^ and plotted on a log10scale. n=3 donors in **a,d-h**; n=2 donors in **b.** Individual donors are represented by unique symbol shapes, while the dark symbols with a black outline represent the donor-specific averages. Bars represent mean and standard deviation. Statistical significance was assessed using two-factor ANOVA with donor and condition as main effects, followed by Tukey’s HSD post hoc test for pairwise comparisons between conditions. Conditions sharing a common letter are not significantly different (P ≥ 0.05); conditions with no shared letters are statistically significant (P < 0.05).

Next, we assessed soluble tissue inhibitor of metalloproteinases-1 (TIMP-1) as a regulator of ECM turnover (Fig. 3d). TIMP-1 showed an inverse trend with sGAGs; PC significantly reduced TIMP-1 compared to controls at d7, and returned to baseline at d14, likely reflecting an adaptive transient response. Together, these complementary measurements demonstrate that cartilage degradation was associated with HPC, whereas PC had a protective effect, preserving matrix integrity.

To assess the overall transcriptomic response to compressive strain, a principal component analysis (PCA) was performed. PCA revealed distinct separation between mechanically loaded and static samples, with Euclidean distances (ED) of 4.48 for HPC and 3.36 for PC compared to static control clusters (Fig. 3e; Extended Data Fig. 1c,d). However, there was substantial overlap between PC and HPC clusters (ED: 1.69), suggesting that varying strain levels did not induce significantly divergent transcriptomic profiles. Concordant with observed proteoglycan loss, HPC significantly upregulated the gene expression of mechanosensitive ion channel, *TRPV4* (Fig. 3f; full gene expression panel, Extended Data Fig.1e-h), and non-significantly increased matrix degrading enzyme, *ADAMTS5 (*Fig. 3g), whereas PC showed no apparent changes compared to the control. HPC and PC both increased *PIEZO1* significantly (Fig. 3f) and *MMP13* non-significantly (Fig. 3g). Cartilage-specific markers *ACAN* and *COL2A1* (Fig. 3h) were both decreased with mechanical loading, but only *ACAN* significantly decreased with HPC. Through PD-CartChip, we demonstrated that the platform integrated with end-stage KOA explants was sensitive to strain at the ECM and protein levels, but showed only marginal differences at the transcriptomic level.

### PD-CartChip model is sensitive to different cartilage degradation stressors

We explored mechanical overloading via HPC and hyperinflammation via supraphysiological levels of pro-inflammatory cytokine stimulation (INF) as examples of known stressors that drive KOA cartilage degradation in our model. PCA on global gene expression profiles revealed a clear separation between the two stressor clusters (ED: 6.78, Fig. 4a; Extended Data Fig. 2a). INF demonstrated the largest shift (ED: 7.40) from baseline, followed by HPC (ED: 2.09). Two-way hierarchical clustering further revealed distinct transcriptomic profiles between HPC and INF-stressed PD-CartChip models (Fig. 4b). *PIEZO1* gene expression was significantly upregulated by HPC and downregulated by INF compared to the control (Fig. 4c; full gene expression panel, Extended Data Fig. 2b-e). INF significantly downregulated cartilage-specific genes (*ACAN, COL2A1)* and upregulated catabolic (*ADAMTS5, MMP13)* and inflammatory genes (*IL6, IL1β*) (Fig. 4d-f). HPC induced a similar but attenuated trend. HPC significantly upregulated lubricin (*PRG4*) and collagen type X (*COL10A1),* and non-significantly increased collagen type I (*COL1A1,* Fig. 4g), demonstrating a compensatory response and matrix remodelling towards fibrocartilage and hypertrophic cartilage.

**Fig. 4.**
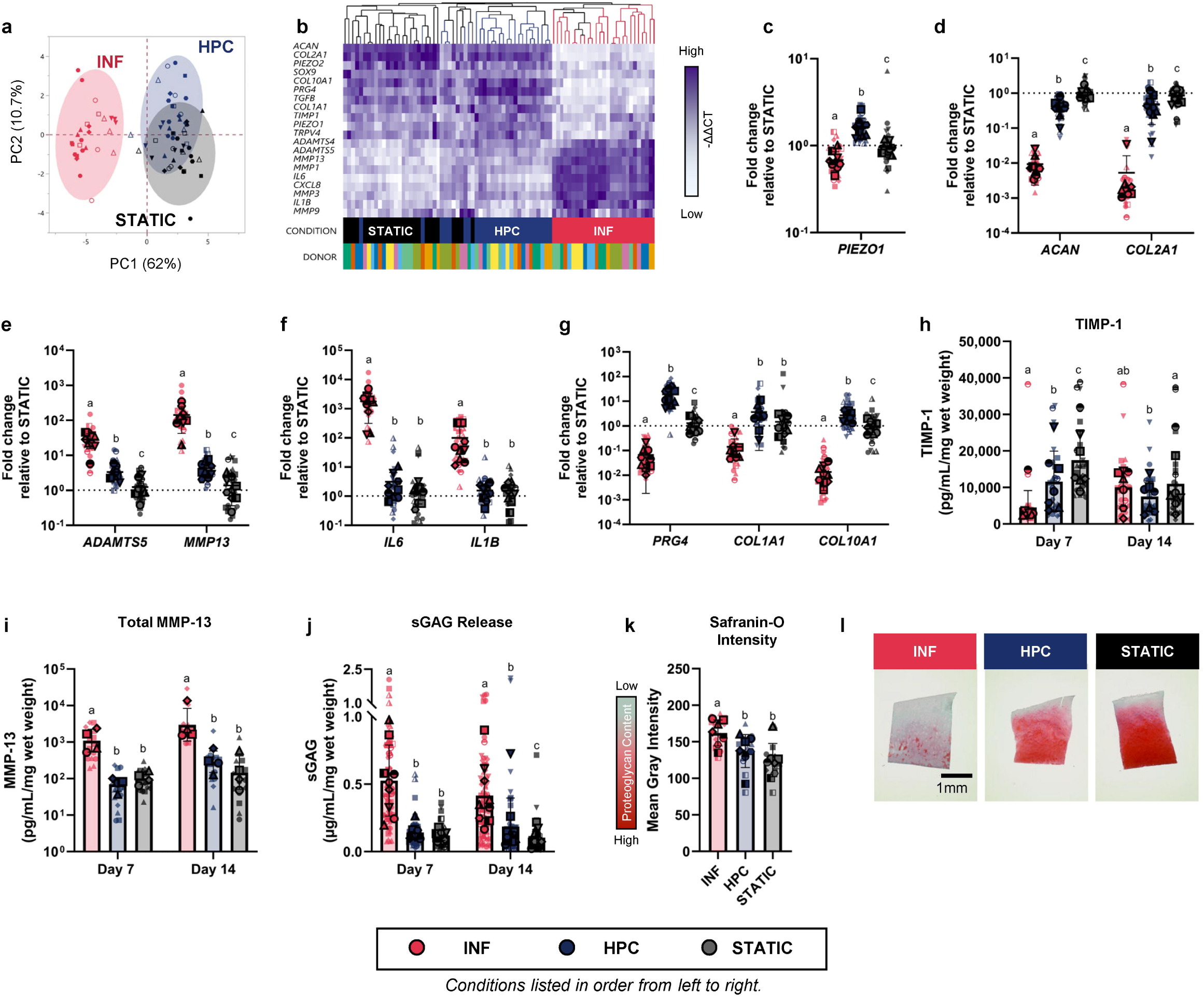
PD-CartChip model is sensitive to mechanical overload (HPC) and hyperinflammatory (INF) stressors. **a,** Principal component analysis (PCA) illustrating 2D clustering of individual samples based on donor-normalized gene expression (-ΔΔCT) across principal component 1 (PC1, 62% variance) and principal component 2 (PC2, 10.7% variance); n=9 donors. Shaded regions represent 95% confidence ellipses for each respective group. **b,** Two-way hierarchical clustering heatmap of donor-normalized gene expression (-ΔΔCT) for individual samples across conditions, grouping replicates by similar gene expression with purple indicating high relative gene expression levels and white indicating low relative levels. Donor-blocked relative fold change in cartilage gene expression at Day 14 across **c,** mechanosensitive ion channels *(PIEZO1),* **d,** cartilage-specific *(ACAN, COL2A1),* **e,** catabolic enzymes (*ADAMTS5, MMP13),* **f,** inflammatory (*IL6, IL1B*), and **g,** matrix remodeling (*PRG4, COL1A1, COL10A1*) markers, calculated via 2^-ΔΔCT^ and plotted on a log10scale. **h,** Soluble tissue inhibitor of matrix metalloproteinases-1 (TIMP-1, pg/mL) levels normalized to initial tissue wet weight (mg). **i,** Total matrix metalloproteinase-13 (MMP-13, pg/mL) normalized to initial tissue wet weight (mg), plotted on a log10scale. **j,** Sulfated glycosaminoglycans (sGAG, µg/mL) release from the tissue to supernatant normalized to initial tissue wet weight (mg). **k,** Quantification of Safranin-O/Fast Green staining by mean gray intensity, where higher values indicate greater proteoglycan loss. **l,** Representative histological images of Safranin-O/Fast-Green stained cartilage sections at Day 14 where intense red staining corresponds to rich proteoglycan content, while a reduction in red intensity indicates matrix degradation. Scalebar = 1mm. n=9 donors in **a-g,j**; n=7 donors in **h,k**; n=4 donors in **i**. Individual donors are represented by unique symbol shapes, while the dark symbols with a black outline represent the donor-specific averages. Bars represent mean and standard deviation. Statistical significance was assessed using two-factor ANOVA with donor and condition as main effects, followed by Tukey’s HSD post hoc test for pairwise comparisons between conditions. Conditions sharing a common letter are not significantly different (P ≥ 0.05); conditions with no shared letters are statistically significant (P < 0.05).

Distinct gene signature profiles were accompanied by stressor-dependent soluble responses that were different in magnitude and temporal dynamics. INF induced a significant, transient decrease in TIMP-1 at d7, whereas HPC significantly reduced TIMP-1 throughout the culture (Fig. 4h). Total matrix metalloproteinase 13 (MMP-13) significantly increased with INF and remained elevated at d14, while HPC showed a non-significant increase at d14 (Fig. 4i). Released sGAGs were significantly increased with INF throughout the culture, whereas HPC showed a delayed response, reaching significantly elevated levels versus control at d14 (Fig. 4j; Extended Data Fig. 2f). This coincided with significant proteoglycan loss with INF and moderate, non-significant loss under HPC at d14 (Fig. 4k,l; Extended Data Fig. 2g). These findings confirmed that the PD-CartChip could capture distinct stressor-dependent responses and demonstrated that while INF and HPC stressors elicited partially distinct molecular and temporal responses, both converged on cartilage degradation.

### Perturbation of stressed PD-CartChip model with dexamethasone

The PD-CartChip models were tested against DEX (100nM), a clinically relevant corticosteroid KOA treatment. PCA of the gene expression profiles revealed a global transcriptional suppression with DEX treatments, characterized by parallel and near identical ED of 3.42 for HPC and DEX (HPC+DEX) and 3.21 for INF and DEX (INF+DEX) relative to respective non-treated, stressor conditions (Fig. 5a; Extended Data Fig. 3a). This was corroborated by a two-way hierarchical clustering analysis that revealed an overall decrease in catabolic and inflammatory gene expression and partially sustained anabolic and cartilage-specific expression with DEX, regardless of stressors (Fig. 5b). There was a subset of genes that exhibited concordant, stressor-independent responses to DEX (full gene expression panel, Extended Data Fig. 3b-e). DEX significantly reduced matrix-degrading enzymes (*ADAMTS5, MMP13)*, interleukin-6 (*IL6)*, and *COL2A1* expression (Fig. 5c-e), and significantly increased *ACAN* expression (Fig. 5e) relative to respective non-treated controls across all stressor conditions. *COL1A1* and *COL10A1* expression significantly decreased with HPC+DEX, whereas reductions with INF+DEX were minimal relative to their respective non-treated controls (Fig. 5f). A distinct subset of genes exhibited stressor-dependent responses to DEX, with distinct expression changes across stressors. Specifically, DEX significantly reduced interleukin-1 beta (*IL1β*) expression under INF, but not HPC conditions (Fig. 5d). *PRG4* expression decreased significantly with HPC+DEX but increased with INF+DEX (Fig. 5f).

**Fig. 5.**
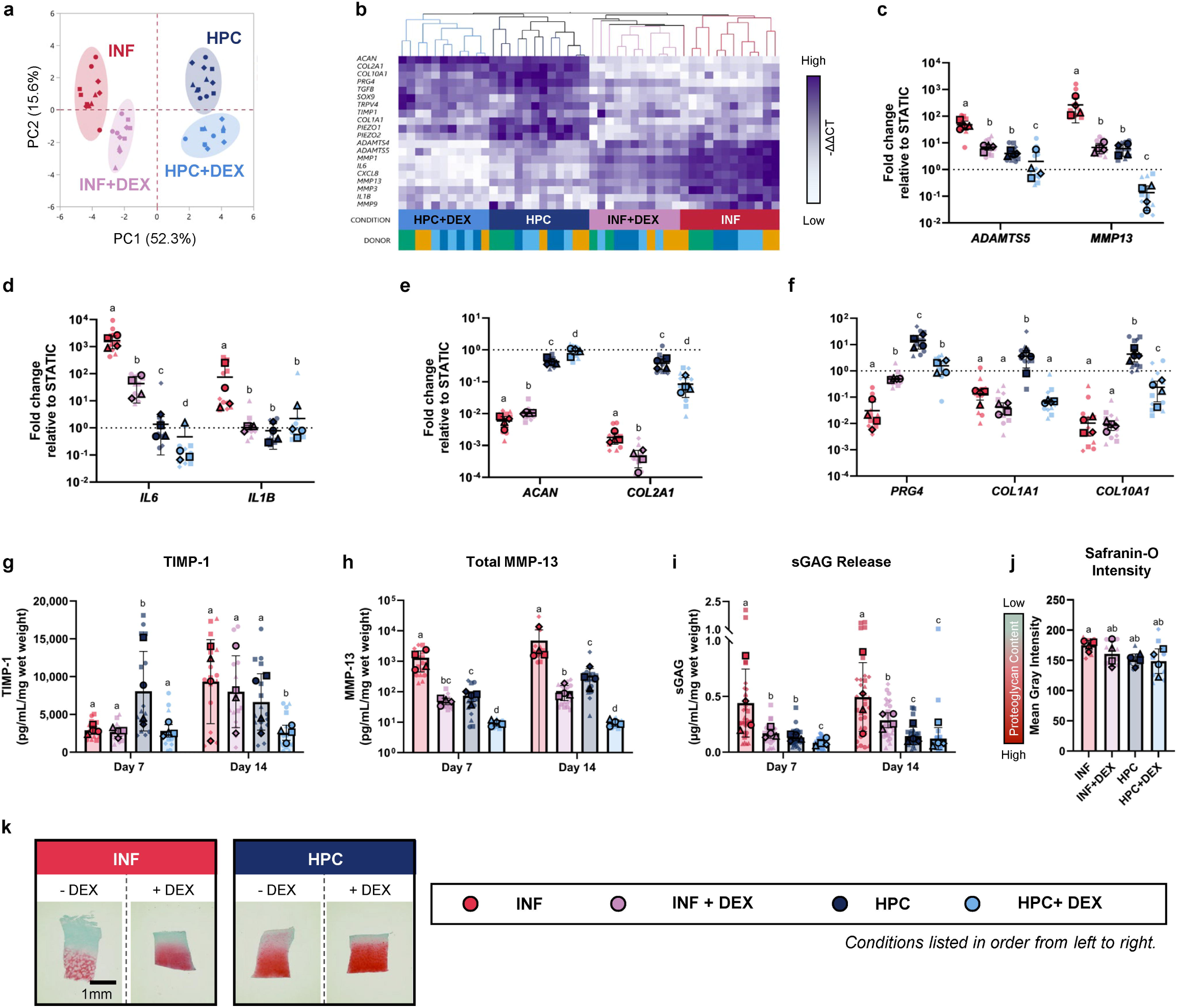
Perturbation of stressed PD-CartChip model with dexamethasone (DEX). **a,** Principal component analysis (PCA) illustrating 2D clustering of individual samples based on donor-normalized gene expression (-ΔΔCT) across principal component 1 (PC1, 52.3% variance) and principal component 2 (PC2, 15.6% variance). Shaded regions represent 95% confidence ellipses for each respective group. **b,** Two-way hierarchical clustering heatmap of donor-normalized gene expression (-ΔΔCT) for individual samples across conditions, grouping replicates by similar gene expression with purple indicating high relative gene expression levels and white indicating low relative levels. Donor-blocked relative fold change in cartilage gene expression at Day 14 across **c,** catabolic enzymes (*ADAMTS5, MMP13),* **d,** inflammatory (*IL6, IL1B*), **e,** cartilage-specific *(ACAN, COL2A1),* and **f,** matrix remodeling (*PRG4, COL1A1, COL10A1*) markers, calculated via 2^-ΔΔCT^ and plotted on a log10scale. **g,** Soluble tissue inhibitor of matrix metalloproteinases-1 (TIMP-1, pg/mL) levels normalized to initial tissue wet weight (mg). **h,** Total matrix metalloproteinase-13 (MMP-13, pg/mL) normalized to initial tissue wet weight (mg), plotted on a log10scale. **i,** Sulfated glycosaminoglycans (sGAG, µg/mL) release from the tissue to supernatant normalized to initial tissue wet weight (mg). **j,** Quantification of Safranin-O/Fast Green staining by mean gray intensity, where higher values indicate greater proteoglycan loss. **k,** Representative histological images of Safranin-O/Fast-Green stained cartilage sections at Day 14 where intense red staining corresponds to rich proteoglycan content, while a reduction in red intensity indicates matrix degradation. Scalebar = 1mm. n=4 donors. Individual donors are represented by unique symbol shapes, while the dark symbols with a black outline represent the donor-specific averages. Bars represent mean and standard deviation. Statistical significance was assessed using two-factor ANOVA with donor and condition as main effects, followed by Tukey’s HSD post hoc test for pairwise comparisons between conditions. Conditions sharing a common letter are not significantly different (P ≥ 0.05); conditions with no shared letters are statistically significant (P < 0.05).

At the soluble level, the PD-CartChip model also demonstrated stressor-dependent responses to DEX. DEX significantly reduced TIMP-1 throughout the culture under HPC, with no apparent changes for INF (Fig. 5g). By contrast, DEX significantly decreased MMP-13 under both stressors throughout the culture (Fig. 5h). This differential response highlights stressor-specific modulation of proteolytic activity underlying matrix degradation. Similarly, DEX significantly reduced sGAGs throughout the culture under INF, whereas the reduction under HPC was transient (Fig. 5i; Extended Data Fig. 3f). Consistent with these changes, DEX modestly attenuated proteoglycan loss across stressors, albeit not significantly (Fig. 5j,k; Extended Data Fig. 3g). Taken together, this demonstrates the ability of PD-CartChip to capture concurrent stress-dependent changes to DEX perturbation.

### Individual patient-level responses to dexamethasone treatment

Exploratory analysis in the PD-CartChip model revealed stressor-specific associations across d14 biological readouts (Fig. 6a). Significant associations were designated as coordinated readout “sets” for subsequent donor-level analysis. For INF, increased MMP-13 was surprisingly associated with less tissue-level proteoglycan loss (Set #1). This relationship was also observed under HPC, suggesting a conserved pattern of cartilage tissue changes across distinct stressor conditions; importantly, the inverse correlation indicates that MMP-13 and tissue proteoglycan depletion may reflect distinct rather than overlapping features of matrix degradation.

**Fig. 6.**
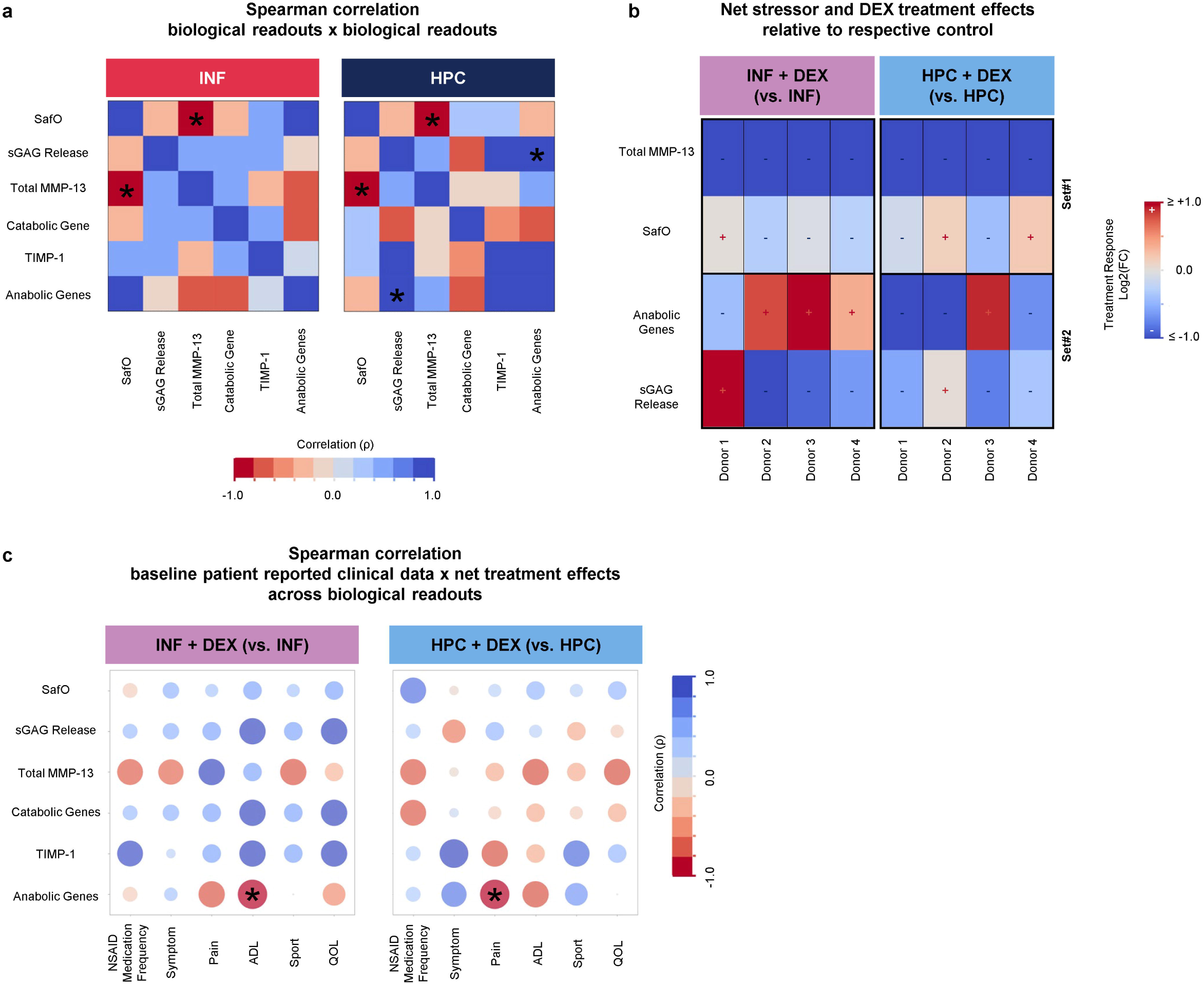
Patient heterogeneity within stress-specific PD-CartChip model. **a,** Non-parametric Spearman rank correlation matrix mapping biological experimental readouts against each other in response to hyperinflammation or mechanical overloading models. Columns represent individual biological experimental readouts at day 14, including Safranin-O (SafO) staining (mean gray intensity), soluble glycosaminoglycans (sGAG Release, µg/mL/mg), tissue inhibitor of metalloproteinases-1 (TIMP-1, pg/mL/mg) and total matrix metalloproteinase-13 (MMP-13, pg/mL/mg), catabolic pathway-related genes (*COL1A1, COL10A1*, *ADAMTS4, ADAMTS5, MMP1, MMP3, MMP9, MMP13, IL6, IL1B, IL8, PIEZO1, PIEZO2;* -ΔCT), and anabolic pathway-related genes (*ACAN, SOX9, COL2A1, PRG4, TGFB, TIMP1, TRPV4;* -ΔCT). The colour gradient scale ranges from strong negative correlation (red, ρ=-1) to strong positive correlation (blue, ρ=+1). **b,** Net stressor and treatment effects relative to non-treated stressed conditions. Values represent log2(fold change relative to the donor-matched reference condition). The colour gradient scale ranges from negative values (<0, blue) representing decreased biological response, to positive values (>0,red) representing increased biological response when treated with DEX. **c,** Non-parametric spearman rank correlation matrix mapping patient clinical history against biological experimental readouts using fold change relative to the donor-matched reference condition. Columns represent patient clinical history including frequent non-steroidal inflammatory (NSAID) intake, and patient-reported outcome measures (PROMs) including, symptoms, pain level, function in activities of daily living (ADL), function in sport and recreation (Sport) and knee-related quality of life (QOL) retrieved from knee injury and osteoarthritis outcome score (KOOS) subscales reported by patients prior to total knee arthroplasty surgery. The raw KOOS subscales were inverted where higher scores denote greater functional impairment and worsening clinical severity. The colour gradient scale ranges from strong negative correlation (red, ρ=-1) to strong positive correlation (blue, ρ=+1) and the size of each bubble corresponds to the absolute magnitude of the correlation to identify key clinical drivers linked to donor variability. n=4 donors. Asterisks indicate significant differences (*P <.05).

By contrast, increased expression of the curated anabolic pathway-related gene set was associated with increased sGAG release under HPC (Set #2); this supports a compensatory anabolic response to matrix loss.

Donor-specific treatment responses were then examined for concordance with the direction of the relationships identified in the untreated stressed models (Fig. 6b). Set #1 was common across stressors, but treatment responses were donor-specific, with decreased MMP-13 and increased proteoglycan loss observed only in Donor 1 under INF. Donors 2 and 4 showed similar patterns, under HPC. Donor 3 showed a distinct response, with decreased MMP-13 and proteoglycan loss, suggesting reduced matrix degradation.

Set #2 was unique to HPC and exhibited donor-specific treatment responses, with concordant responses limited to Donors 1 and 4, characterized by decreased expression in curated anabolic pathway-related genes and sGAGs. Donors 2 and 3 under HPC showed discordant responses, and the pattern was absent across all donors under INF.

Together, these findings demonstrate that stressor-associated biological relationships were not uniformly preserved following DEX treatment, revealing donor-dependent response patterns.

Next, as part of an exploratory analysis to highlight the model’s potential to contextualize clinical correlates, we assessed associations between stressor-specific treatment responses and baseline donor clinical features across different stressors (Fig. 6c). Attenuated treatment-induced anabolic gene response was strongly associated with poorer patient-reported daily function at baseline in the INF group and with poorer patient-reported pain at baseline in the HPC group. Together, this suggests that donor clinical history may influence treatment sensitivity in a stressor-dependent manner. These exploratory findings demonstrate the PD-CartChip model’s sensitivity and potential utility for integrating patient clinical annotations.

## Discussion

In alignment with recent global regulatory shifts towards new approach methodologies (NAMs), we present a patient-specific COC model, PD-CartChip, that provides a testing platform for understanding KOA pathogenesis and evaluating novel treatments while incorporating patient heterogeneity. Our model uniquely leverages end-stage KOA patient-derived articular cartilage explants, exposed to disease-relevant stressors including mechanical overloading via controlled, unconfined compression or hyperinflammation via pro-inflammatory cytokine stimuli, and captures distinct stressor- and patient-donor associated response profiles to corticosteroid treatments.

PD-CartChip captured baseline responses induced by established KOA-relevant stressors, mechanical overload and hyperinflammation, and revealed tissue profiles that putatively reflect emerging OA biochemical signatures of OA endotypes^5,17,18^. For example, hyperinflammation and excessive loading elicited different severities of proteoglycan loss alongside distinct temporal patterns of proteolytic regulation. Mechanical overloading also triggered an inflammatory transcriptional response, congruent with mechanoinflammation^19^, albeit attenuated relative to hyperinflammatory stressor responses, highlighting overlapping biological processes across distinct stressors. This overlap parallels KOA patients who present with clinically similar OA features despite differing underlying etiologies. The PD-CartChip model provides an opportunity to study these distinct yet interconnected disease processes and how they shape degradative trajectories.

This complexity of KOA disease trajectories underscores the value of human end-stage KOA explants as a biologically relevant model system. Although select KOA features can be recapitulated in existing cell-hydrogel based COC models^9,12,13^, KOA explants preserve native tissue organization and cell-matrix interactions, maintaining a pre-existing catabolic milieu and donor-specific responsiveness that provides the necessary complex backdrop for evaluation of therapeutics. To illustrate, *PRG4* gene expression was upregulated with mechanical overloading in PD-CartChip, contrary to previous studies reporting its downregulation^13^. This conflicting *PRG4* response may partly reflect differences in native zonal and matrix content^20^ in diseased cartilage versus cell-encapsulated hydrogels. While PRG4, a normally abundant protein in cartilage, is often reduced in KOA^21–23^, intact superficial zone chondrocytes can upregulate *PRG4* transcription as an adaptive response to preserve tissue function^24^. The upregulation in our PD-CartChip reflects the capacity to mount such adaptive responses.

Cell-hydrogel constructs, by comparison, commonly use dedifferentiated primary human chondrocyte culture involving 2D expansion, thereby inducing selection bias, and re-differentiation into uniform 3D constructs that collectively result in reduced native heterogeneity in chondrocyte populations and their corresponding spatial cell distribution^25,26^. Generally, they do not fully recapitulate the complex zonal and spatial matrix organization of native cartilage. Additionally, while advantageous for specific controlled studies of chondrocyte behavior and highly scalable, these constructs are substantially less stiff compared to native human cartilage^27^. Marked differences in mechanical stiffness between explants and cell-hydrogel constructs create challenges for sample dimensional variation, scaling, and translational applicability. Several cell-hydrogel COC platforms rely on confined compression. This test configuration is valuable for capturing the biphasic, fluid-pressurization-dependent behavior of native cartilage^28,29^. However, they reveal stress-strain relationships, fluid pressurization, and load transmission that fall short of endogenous mechanical behavior^30^. Geometrically constrained, double-network, or decellularized-ECM-based constructs better approximate native stress-strain profiles and equilibrium moduli, though direct evidence of fluid pressurization and cell-scale load transmission in these systems remains limited^31,32^.

Moreover, unlike uniform constructs that can be simultaneously set to a single strain target across multiple samples for high-throughput use^33^, explants vary significantly in sample thickness due to anatomical site-specific differences^34,35^. Accommodating this variability requires several individual actuators to deliver cyclic loading regimes, consequently increasing device complexity and limiting throughput. PD-CartChip resolves this bottleneck by utilizing acrylic flexures and manual adjustments to set strain targets per sample and employing a single actuator across multiple samples and devices. Accordingly, the number of devices per single actuation platform is limited by the linear actuator load capacity and available stacking height. Currently, PD-CartChip accommodates two devices per platform, but it can be scaled for future studies requiring higher throughput. Integrating this platform with explant tissue enables mechanically driven biological interrogation of clinically relevant tissue and true disease-modifying capability of therapeutic treatments.

Consistent with this, we showed that PD-CartChip is sensitive to biological shifts in stressor-specific responses to DEX, a potent anti-inflammatory treatment^36^ known to provide short-term KOA symptom relief. Despite its anti-inflammatory action, DEX ameliorated catabolic responses induced by mechanical overload and hyperinflammation, while also suppressing pro-anabolic *COL2A1* expression. These competing responses to DEX align with both anti-catabolic and anti-anabolic reported effects sustained following repeated clinical use^37,38^.

Although the net therapeutic effect of DEX reflected an overall improvement in cartilage degradation, donor-specific modulation was apparent, showing different concordant and discordant changes across the multivariable model readouts. This prompted exploratory stressor-and donor-level analyses of coordinated changes in DEX, highlighting the sensitivity of the PD-CartChip model to capture identifiable associations within our small cohort of donors. Despite the small cohort, we observed substantial donor-level variation, even within the same stressor context, with only some donors exhibiting concordant directional responses. These donor-level patterns may help define subcategories of treatment responsiveness. Exploratory correlations also revealed distinct sets of coordinated changes that were either common or distinct across untreated, stressed conditions. In the future, broader sets of such relationships could be used to assist in identifying signatures associated with underlying stressor conditions, analogous to putative molecular endotypes emerging based on biochemical analysis^5,17,18^. Future applications of PD-CartChip could characterize inherent patient-specific signatures that may reflect diverse OA endotypes without external stressors and be used to probe endotype-associated therapeutic responses.

Model readouts were contextualized using baseline patient clinical features. Patients with worse baseline daily function and pain showed lower anabolic gene responses following DEX treatment under hyperinflammatory and mechanical overloading stressors, respectively. These findings suggest attenuated DEX treatment responsiveness in donors with poorer baseline patient-reported outcomes, mirroring clinical observations linking persistent DEX use with diminished anabolic benefits and adverse effects^37,38^. Integrating clinical annotations therefore enables a more holistic interpretation of donor-specific treatment responses.

The PD-CartChip shows promise for future applications but remains subject to important limitations. Longitudinal measurements captured broad temporal dynamics; however, correlation analyses were limited to time-matched endpoint readouts, potentially missing early or transient relationships. We used a curated subset of biological readouts, but expansion to omics-based readouts could capture unbiased changes. This could be combined with machine learning to provide broader disease-associated and treatment-responsive signatures for donor and stressors, aligned with other profiling approaches^5,17,39^, and current NAM principles.

Site-specific anatomical harvest sites in the cartilage were not accounted for and may present an additional source of heterogeneity in our results^40^. Explants from multiple sites within the same donor have previously been treated as independent samples^41^ due to limited tissue availability. Here, explants were treated as independent samples with donor included as a model effect to account for samples originating from the same individual. Multiple mechanical factors act on the knee joint, including compression, shear stress, and interstitial fluid pressure^42^, whereas our model was limited to compressive forces. Uncontrolled shear stresses were also likely produced due to frictional contact between the paddle and explant, geometric well constraints, and tissue heterogeneity, and they will be considered in future iterations in the model. The subchondral bone is also an important contributor to mechanotransduction^43^ but was omitted from this study to isolate cartilage-specific responses and eliminate confounding effects. Bone and other joint tissues can be incorporated in future iterations of the PD-CartChip in a modular fashion, as we have done with static KOA^44^ and inflammatory arthritis models^45^, and as demonstrated by other groups^12,13^.

## Conclusion

We successfully developed a novel patient-specific, PD-CartChip, that integrated KOA end-stage cartilage explant tissue with disease-relevant stressors, including mechanical overloading and hyperinflammation. Multivariable biological readouts captured stressor-dependent and donor-specific responses to an anti-inflammatory treatment. Linking these responses with clinical annotations provided patient-specific context and established a framework that can be expanded in future studies to investigate unique OA mechanisms, aid in patient stratification, or evaluate the performance of the latest DMOADs under defined joint stressors.

## Materials and Methods

### PD-CartChip device design and fabrication

The completed PD-CartChip platform was designed to apply controlled compressive loading to cartilage explants housed within individual microwells. Compression was achieved through deformation of compliant cantilever flexures coupled to an Arduino-controlled servo motor, enabling programmable loading regimes. The displacement range of the carriage assembly was constrained using adjustable limit and stop screws, permitting precise control over tissue compression depth.

The PD-CartChip was designed in-house and fabricated primarily from ultraviolet (UV)-scratch-resistant cast poly(methyl methacrylate) (PMMA) sheets (McMaster-Carr), together with several custom 3D-printed components. All PMMA components were manufactured using a computer numerical control (CNC) micromilling machine (Datron Neo, Datron Dynamics)^46^. The device consisted of three principal subsystems: (i) a microwell chip layer for tissue culture, (ii) a compliant flexure spring layer containing the compression paddles and flexure beams, and (iii) a carriage frame used for actuation and displacement control. Each microwell chip layer consists of 12 total microwells, four of which can be exposed to mechanical loading and the other eight for static explant culture.

Each layer of the flexural assembly was machined individually from PMMA sheets prior to assembly. Structural components including the flexure layer, paddles, and carriage frame were bonded using cyanoacrylate adhesive (Loctite 401), whereas fluidic components including the microwell plate and microwell base were solvent bonded to achieve a uniform, optically clear, and leak-resistant seal.

For solvent bonding, the mating PMMA surfaces were first cleaned thoroughly, after which 99% ethanol was evenly applied to the bonding interface. Components were aligned and compressed at approximately 1000 lbs for 3 min at 70°C using a heated hydraulic press (Carver, Inc.)^47,48^. Following bonding, the flexure layer and carriage frame were positioned above the microwell assembly and aligned using stainless steel dowel pins. Compression paddles were inserted into precision-machined notches within the flexure layer and aligned concentrically with the underlying microwells prior to adhesive fixation.

Additional enclosure components, including the external reservoir casing and lid assembly, were fabricated using CNC micromilling. The transparent lid pane and lid frame were solvent bonded using the same thermal-assisted bonding protocol described above.

To improve optical clarity and surface finish, all PMMA surfaces subjected to full surface-facing operations were vapor-polished using dichloromethane (DCM; ≥99.8%, Sigma-Aldrich). Prior to polishing, surfaces were cleaned using compressed air followed by 70% ethanol. Approximately 20–30 mL of DCM was heated within a sealed Pyrex flask on a hotplate inside a chemical fume hood until vapor was generated. PMMA components were then briefly exposed to the solvent vapor for 3–5 seconds at approximately 5 mm from the flask opening. Polished components were allowed to dry for at least 3 hr prior to cleaning and device assembly.

Actuation components and modular mounting brackets were fabricated using fused deposition modeling (FDM) 3D printing on a Bambu X1 Carbon printer (Bambu Labs) using carbon fiber-reinforced PA6 nylon filament (Polymaker). Components were printed using seven perimeter walls and 35% gyroid infill to improve structural rigidity. Brass threaded inserts were incorporated into all mechanically loaded fastening interfaces.

Prior to each experiment, devices were sterilized using 10% bleach followed by 70% ethanol and allowed to air-dry overnight inside a biosafety cabinet. Microwells were rinsed three times with phosphate-buffered saline (PBS) and surrounding reservoir chambers were filled with PBS to minimize evaporative losses during long-term culture experiments.

### Cartilage characterization, device optimization, and endpoint design

The PD-CartChip device design was optimized specifically towards supporting the load requirements of the native cartilage samples such that a sufficient travel range and stiffness level was achieved to bring the samples to target 2 mm diameter native cartilage samples ranging from 1-3 mm in length were mechanically characterized using the Dynamic Mechanical Analyzer (DMA) Q800 (TA Instruments) to determine the average peak load limit required for the internal flexure array (Supplementary Fig. 2a, 3a) and subsequently tested under sinusoidal loading using a custom heated water bath design integrated into the UniVert mechanical testing system (CellScale; Supplementary Fig. 2b, 3b). Using this load limit, both linear and non-linear beam bending theories were used to eliminate any extraneous design alternatives (Supplementary Fig. 4, 5). The final design configuration was then assessed for actual loading responses, and motion accuracy using a Instron 5848 MicroTester (Instron Inc.; Supplementary Fig. 6, 7) and Keyence VHX 7000 (Keyence Inc.) digital microscope (Supplementary Fig. 8), respectively. Details of the comprehensive design optimization pipeline can be found in the Supplementary Notes.

The final design derived from this optimization procedure was the PD-CartChip platform which allows for multiplexed controlled unconfined compression for native cartilage explant logs up to 2.5 mm in thickness (Supplementary Fig. 1). The device is comprised of a system-level assembly and a device-level assembly of components. At the device level, 2 mm cartilage explants are placed along their length into a 200 µl microwell. The microwell maintains multiple 500 µm and 1 mm wide side channels to allow for fluid movement in and out of the explants under compression to mimic an unconfined configuration (Fig. 1b(ii)). In addition, this well geometry maintains clearance on either side to accommodate lateral expansion of the explants. Acrylic paddles are used to facilitate the contact and load application to the explants by delving into the microwells vertically. The multiplexing capabilities of the system are made possible using compliant acrylic flexures which facilitate independent strain targeting of explants through the manual adjustment of limit screws depicted in (Fig. 1b(i)). At the device level there are two types of flexural arrays: the internal flexures and the external flexures (Fig. 1b(i)). The internal flexures support the paddle in contact with the cartilage and are translated through mounts directly attached to the actuator. The external flexures ensure that the frame moves linearly along the path of actuation. The paddle and internal flexures go through three primary phases to facilitate independent compression. (1)

Phase 1, uncompressed phase: the paddle and internal flexures start at rest prior to cycle start (Fig. 1c(i)). (2) Phase 2, dynamic compression phase: once the cycle starts the sample is compressed at a frequency of 1Hz to the compressive strain target (10% for physiological strain, 30% for hyperphysiological strain). As this is happening the internal flexures act as resistive spring elements to ensure that the paddle reaches its compression target (i.e., the strain maximum) without the cartilage deflecting the paddle position (Fig. 1c(ii)). (3) Phase 3, hold phase: once at target the paddle is stopped by a set limit screw, thus holding the position for that particular sample. The frame continues its linear motion to compress replicates mounted elsewhere in the array (Fig. 1c(iii)).

The key design focus of the optimization procedure was maintaining enough stiffness for the internal array to reasonably resist the peak load generated by the cartilage samples and offering enough compliance to allow for varying sample lengths of explants. The flexures selected were chosen to accommodate the peak loads taken from off-device cartilage testing, while maintaining a workable footprint for the assembly on a microscope and incubator. Zooming out from the device level to the system assembly, the entire internal device assembly is situated in an acrylic casing to prevent humidity loss and maintain sterility. The casing is secured to an acrylic mounting plate using 3D printed mounting brackets which holds the case in compression from either end (Fig. 1a). The actuator positioned in front of the casing utilizes a 75 kg servo coupled with a rack and pinion system to drive the internal device flexures using a single connecting rod per device (Fig. 1a). Up to three devices can be accommodated per assembly and placed on a single rack in a standard incubator.

### Human cartilage sample collection and explant culture

Human articular cartilage from end-stage OA (Kellgren-Lawrence Grade 3-4) total knee arthroplasties were acquired with patient consent and institutional ethics approval (UHN REB#14-7483). Patient information (n=9 donors), including sex, age, and body mass index (BMI), are provided in Supplementary Table 1. Full-depth cartilage was removed from the medial femoral condyle and sectioned with a punch biopsy (Integra Miltex, 2 mm diameter). Explants were incubated in a 96-well plate for a 3- to 5-day acclimatization period as before^44^, prior to starting experiments.

Optimized explant medium is a 1:1 ratio of Roswell Park Memorial Institute (RPMI) 1640 and Dulbecco’s modified Eagle medium (DMEM), low glucose, supplemented with 1X insulin-transferrin-selenium, 1 mM sodium pyruvate, 50 μg/mL L-ascorbic acid 2-phosphate, 50 μg/mL L-proline, 1% gentamicin^44^.

### Cartilage degradation induction and perturbation in the PD-CartChip model

To generate a mechanical overload model, cartilage explants were subjected to 30% unconfined compressive strain (HPC). Mechanical loading associated with daily walking was approximated by stimulating samples for 3 hr at 1Hz followed by a 21 hr rest period. This 24 hr loading/rest regimen was based on established cartilage mechanobiology protocols^49^ and is consistent with daily loading/rest regimens previously applied to human osteoarthritis cartilage explants. To generate a hyperinflammatory model (INF), explants were stimulated with a supraphysiological dose of interleukin-1 beta (IL-1β, 5ng/mL, PeproTech) and oncostatin M (OSM, 5ng/mL, PeproTech) known to elicit pro-inflammatory and pro-catabolic effects in static culture^44^. Unstimulated static explants were used as a control group. To evaluate the model under therapeutic perturbation, the optimized culture medium was supplemented with 100 nM dexamethasone (DEX, Bioreagent). This dosage was selected based on a previously established joint-on-a-dish model^44^. The medium was replenished and collected on days 3, 7, and 10. On day 14, both medium and tissue samples were harvested by snap freezing in liquid nitrogen and stored at −80°C for analysis.

### Real-time quantitative polymerase chain reaction (RT-qPCR)

Frozen explant tissue was pulverized using a liquid nitrogen pre-chilled BioPulverizer (BioSpec) for RNA isolation using RNeasy Plus Universal Mini Kit (Qiagen). Two explants were pooled together during processing. cDNA was generated with SuperScript™ IV VILO™ Master Mix (Thermo Fisher). A gene sub-panel was selected based on cartilage-specific genes and mechanoresponsive profiles reported in literature^9,44^. RT-qPCR was run on cartilage-specific gene panels using custom primers (ThermoFisher; Supplementary Table 2) and Taq Pro Universal SYBR qPCR Master Mix (Vazyme) on a QuantStudio™ 5 machine (ThermoFisher). Results were normalized (-ΔΔCT) against reference genes *B2M*, *ACTB*, and *GAPDH* (geometric mean), then to donor-matched STATIC condition median, and plotted as block-centered log10(fold-change=2^-^ΔΔCT).

### Proteoglycan loss, soluble protease, and soluble protein activity

Sulfated glycosaminoglycan (sGAG) content in the conditioned medium was detected by dimethylmethylene blue assay (DMMB) against chondroitin sulfate (sodium salt from shark cartilage) standard A525nm curve. Soluble TIMP-1 and total MMP-13 was detected by Human DuoSet ELISA Development Kits (R&D Systems). All kits were used according to the manufacturer’s protocol. Conditioned medium from two replicates were pooled together for the TIMP-1 and MMP-13 assays. Values below and above assay detection limits were imputed as half the lower and double the upper detection limits, respectively, before applying dilution factors. All soluble factor concentrations were normalized to day 0 cartilage explant wet weight (mg) to account for different-sized cartilage pieces.

### Histology and intensity quantification

Cartilage explants were fixed in 10% neutral buffered formalin for 48 hr at room temperature. The samples were then washed in PBS for 1 hr, dehydrated in anhydrous ethanol for 1 min, and cleared in JFC Solution (Milestone Medical) overnight. The dehydrated samples were embedded in paraffin and sectioned at 5 µM with a manual microtome. Sections were dewaxed with xylene, rehydrated in a series of decreasing ethanol concentrations, and stained with 0.1% (w/v) Safranin-O/0.01% (w/v) Fast Green FCF. The mean gray intensity value from Safranin-O stained sections were quantified in FIJI^50^.

### Statistical analysis

All biological experimental data analyses were performed using JMP®, Version 19 (JMP Statistical Discovery LLC, Cary, NC, 2026). Statistical significance (P<0.05) was assessed using two-factor ANOVA with donor and condition as main effects, followed by Tukey’s HSD post hoc test for pairwise comparisons between conditions. ANOVA assumptions were verified using residuals and Shapiro-Wilk test. Principal component analysis (PCA) and two-way hierarchical clustering were performed on blocked donor-centered -ΔΔCT gene expression values. Correlations were performed using non-parametric Spearman’s rank-order correlation (ρ), and the p-values were adjusted using the Benjamini-Hochberg False Discovery Rate (FDR) procedure. Correlations between biological readouts within stressor-specific models were performed using raw values, including Safranin-O (mean gray intensity), sGAG release (µg/mL/mg), TIMP-1 (pg/mL/mg), total MMP-13 (pg/mL/mg) and gene expression (-ΔCT). Correlations involving treatment responses and patient-reported clinical variables were performed using log_2_fold-change (log_2_FC) values relative to the corresponding stressor only condition. Graphs were generated with GraphPad Prism (version 11.0, GraphPad Software). Detailed statistical results, including P values for group comparisons and correlation analyses, are provided in Supplementary Tables 3 to 7.

### Data sharing

All data generated from this study are available from the corresponding authors upon reasonable request.

## Acknowledgements

We thank Christina Ward and Kim Perry at the Schroeder Arthritis Institute Osteoarthritis Biobank and the Orthopedic Surgery Team and Clinical Research Team for assistance with the consent and acquisition of donor samples. This work is in part supported by the Schroeder Arthritis Institute via the Toronto General and Western Hospital Foundation (University Health Network). This research is funded by the Center for Research and Application of Fluidic Technologies (CRAFT) Project Award, Arthritis Society Canada (Grant#22-0000000123), and Natural Sciences and Engineering Research Council of Canada (NSERC) Discovery Grants to SV (RGPIN-2024-04118, sRGPIN 2018-05737) and EY (RGPIN-2025-06905, RGPIN 2019-05885). We thank CRAFT for access to the Device Foundry at the University of Toronto and technical support. We thank the Advanced Optical Microscopy Facility at the University Health Network. We thank Julie Audet for their guidance in the statistical analysis.

## Author Contributions

L.B., E.Y., S.V., K.P., K.L., and B.M. contributed to the conceptualization of the study. L.B., K.P., and K.C. contributed to the development of the study methodology. R.G. provided human articular cartilage specimens. L.B. performed the biological experiments, with support from B.L., K.W., and K.P. for device set up and operation. K.P., R.C., and E.B. performed the device-characterization experiments. L.B. performed the formal analysis of the experimental data and prepared the associated visualizations. K.P. performed device-related characterization and simulations, and associated analyses and visualizations. L.B. managed and executed the data curation. L.B., S.V., E.Y., and K.P. wrote the original draft of the manuscript. L.B., S.V., and E.Y. reviewed and edited the manuscript. E.Y. and S.V. supervised the project and led funding acquisition. L.B., E.Y., and S.V. were responsible for project administration. All authors reviewers and approved the final manuscript.

