## Extended Data Figures for "Human knee osteoarthritis patient-specific cartilage-on-a-chip model captures donor differences to stressors and treatments"

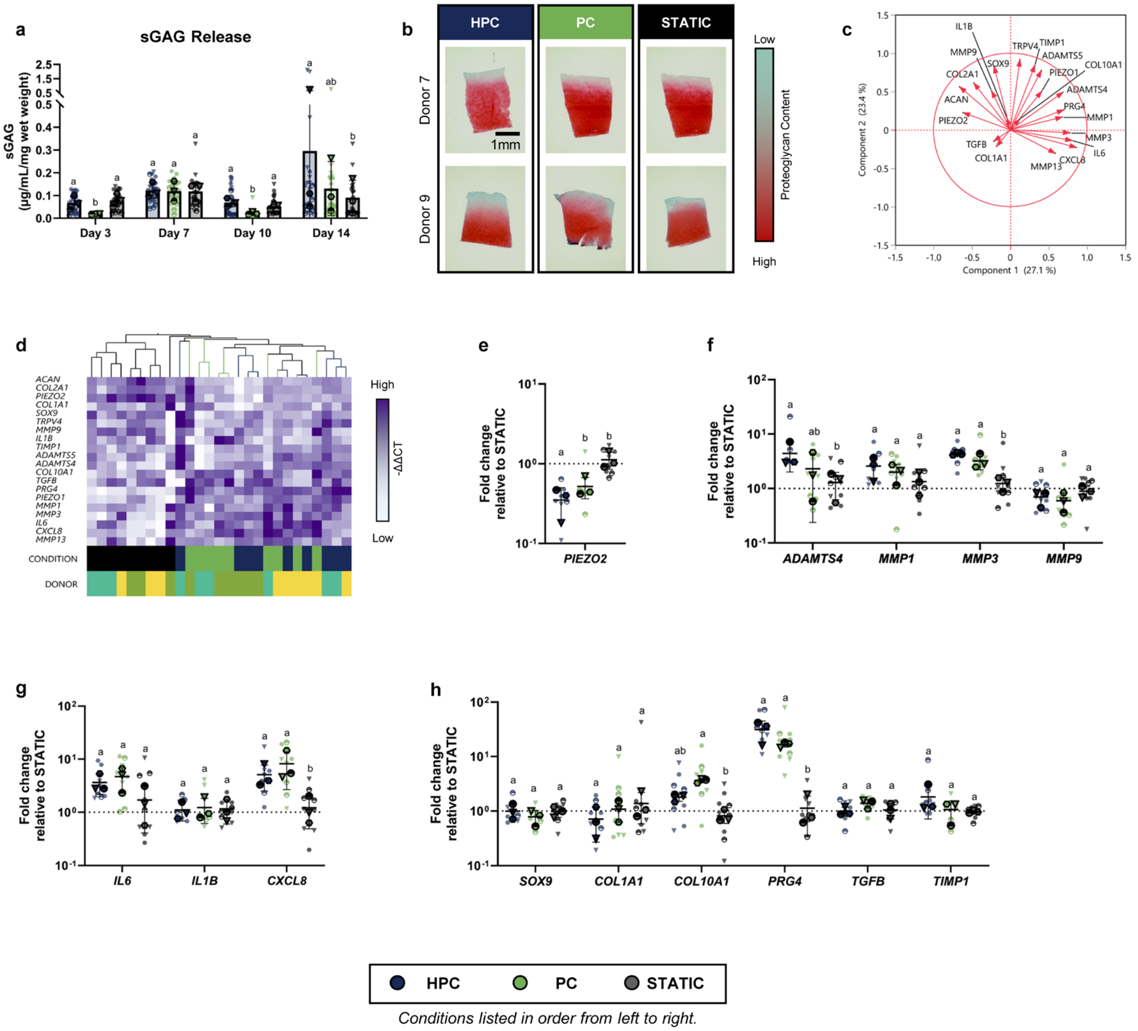


**Extended Data Fig. 1 | Comprehensive dataset for physiological (PC) vs. hyperphysiological (HPC) mechanical loading experiment. a,** Sulfated glycosaminoglycans (sGAG, µg/mL) release from the tissue to supernatant normalized to initial tissue wet weight (mg). **b,** Histological images of Safranin-O/Fast-Green stained cartilage sections at Day 14 where intense red staining corresponds to rich proteoglycan content, while a reduction in red intensity indicates matrix degradation. Scalebar = 1mm. **c,** Principal component analysis (PCA) loading vector plot featuring individual gene contributions to variance based on donor-normalized gene expression (-ΔΔCT) across principal component 1 (PC1, 27.1% variance) and principal component 2 (PC2, 23.4% variance). **d,** Two-way hierarchical clustering of gene expression data set based on donor-normalized gene expression (-ΔΔCT). Donor-blocked relative fold change in cartilage gene expression at Day 14 across **e,** mechanosensitive ion channels *(PIEZO2),* **f,** catabolic enzymes (*ADAMTS4, MMP1, MMP3, MMP9*), **g,** inflammatory (*IL6, IL1B, CXCL8),* and **h,** anabolic and matrix remodeling *(SOX9, COL1A1, COL10A1, PRG4, TGFB, TIMP1)* markers, calculated via $2^{-\Delta\Delta CT}$ and plotted on a log10scale. n=3 donors in **a, c-h**; n=2 donors in **b**. Individual donors are represented by different symbols (symbols with black outline denote donor-specific average).

Statistical analysis was performed using ANOVA with donor and condition as random effects, followed by Tukey's HSD post-hoc test for all pairwise comparisons. Bars represent mean and standard deviation. Conditions sharing a common letter are not significantly different ($P \geq 0.05$); conditions with different letters indicate significant differences ($P < 0.05$).

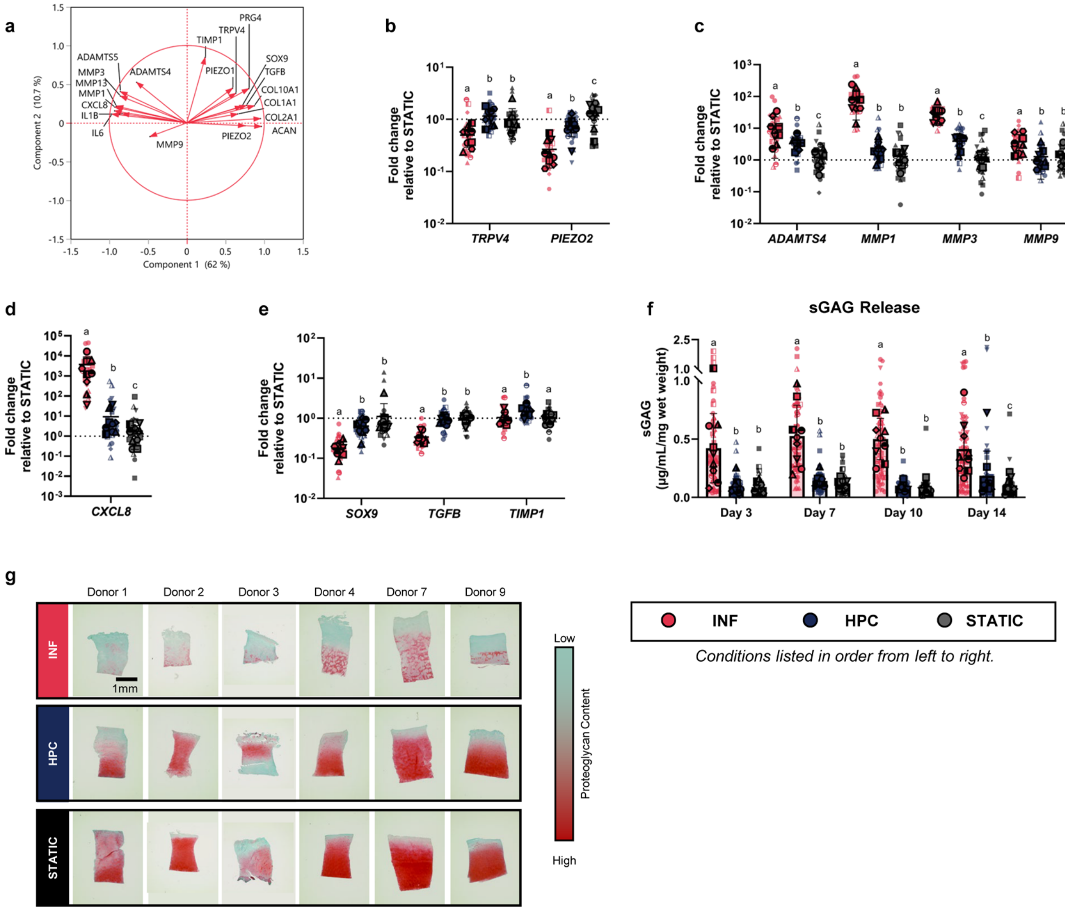


**Extended Data Fig. 2 | Comprehensive dataset for hyperinflammatory (INF) vs. mechanical overload (HPC)-stressed PD-CartChip experiment. a,** Principal component analysis (PCA) loading vector plot featuring individual gene contributions to variance based on donor-normalized gene expression (-ΔΔCT) across principal component 1 (PC1, 62% variance) and principal component 2 (PC2, 10.7% variance). Donor-blocked relative fold change in cartilage gene expression at Day 14 across **b,** mechanosensitive ion channels *(TRPV4, PIEZO2),* **c,** catabolic enzymes (*ADAMTS4, MMP1, MMP3, MMP9*), **d,** inflammatory (*CXCL8)* and **e,** anabolic *(SOX9, TGFB, TIMP1)* markers, calculated via $2^{-\Delta\Delta CT}$ and plotted on a $\log_{10}$scale. **f,** Sulfated glycosaminoglycans (sGAG, µg/mL) release from the tissue to supernatant normalized to initial tissue wet weight (mg). **g,** Histological images of Safranin-O/Fast-Green stained cartilage sections at Day 14 where intense red staining corresponds to rich proteoglycan content, while a reduction in red intensity indicates matrix degradation. Scalebar = 1mm. n=9 donors in **a-f**; n=7 donors in **g.** Individual donors are represented by unique symbol shapes, while the dark symbols with a black outline represent the donor-specific averages. Bars represent mean and standard deviation. Statistical significance was assessed using two-factor ANOVA with donor and condition as main effects, followed by Tukey's HSD post hoc test for pairwise comparisons between conditions. Conditions sharing a common letter are not significantly different ($P \geq 0.05$); conditions with no shared letters are statistically significant ($P < 0.05$).

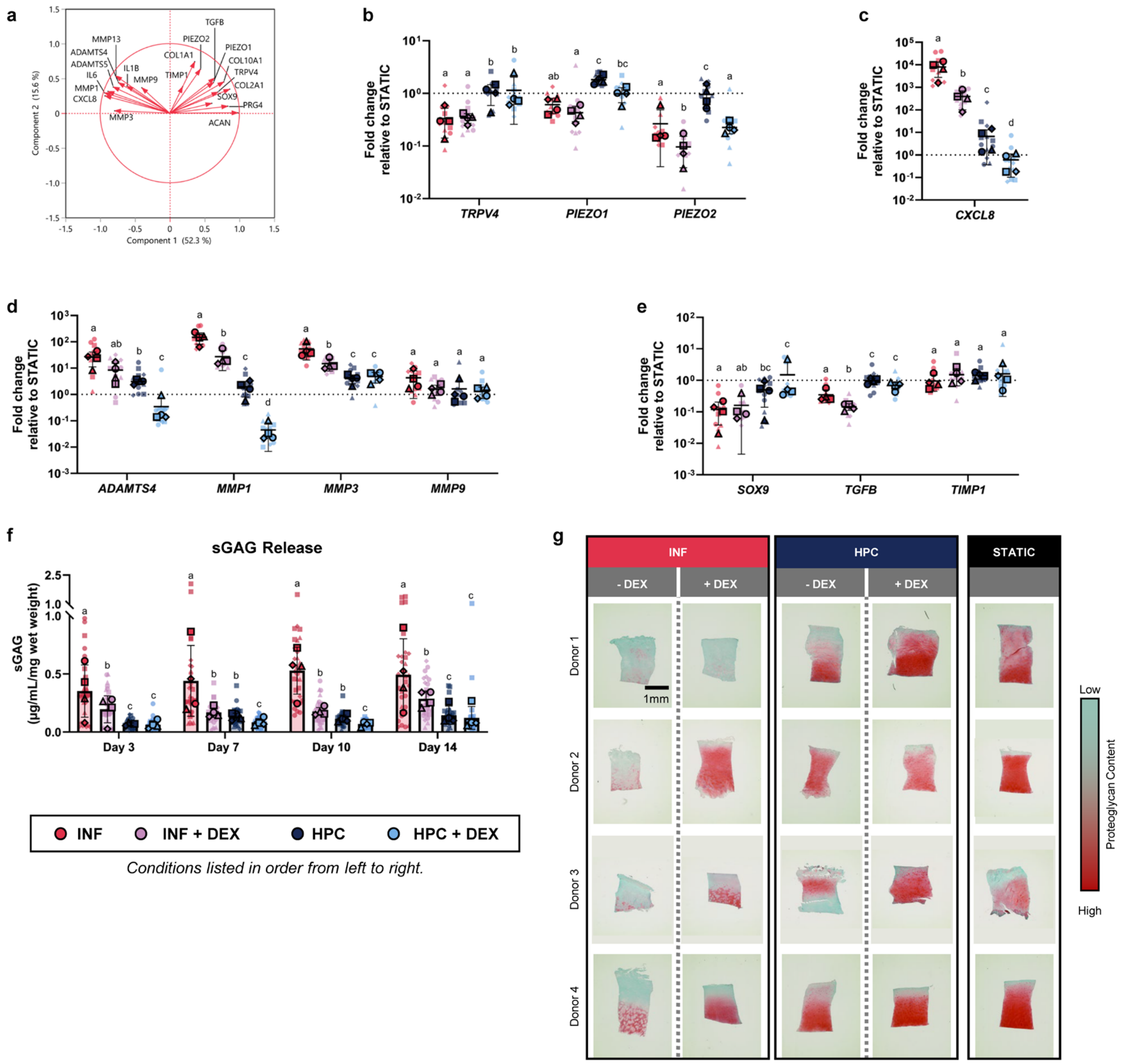


**Extended Data Fig. 3 | Comprehensive dataset for hyperinflammatory (INF) vs. mechanical overload (HPC)-stressed PD-CartChip experiment with dexamethasone (DEX) treatment.** **a,** Principal component analysis (PCA) loading vector plot featuring individual gene contributions to variance based on donor-normalized gene expression (-ΔΔCT) across principal component 1 (PC1, 52.3% variance) and principal component 2 (PC2, 15.6% variance). Donor-blocked relative fold change in cartilage gene expression at Day 14 across **b,** mechanosensitive ion channels *(TRPV4, PIEZO1, PIEZO2),* **c,** inflammatory (*CXCL8),* **d,** catabolic enzymes (*ADAMTS4, MMP1, MMP3, MMP9)* and **e,** anabolic *(SOX9, TGFB, TIMP1)* markers, calculated via $2^{-\Delta\Delta CT}$ and plotted on a log10scale. **f,** Sulfated glycosaminoglycans (sGAG, µg/mL) release from the tissue to supernatant normalized to initial tissue wet weight (mg). **g,** Histological images of Safranin-O/Fast-Green stained cartilage sections at Day 14 where intense red staining corresponds to rich proteoglycan content, while a reduction in red intensity indicates matrix degradation. Scalebar = 1mm. n=4 donors. Individual donors are represented by unique symbol shapes, while the dark symbols with a black outline represent the donor-specific averages. Bars represent mean and standard deviation. Statistical significance was assessed using two-factor ANOVA with donor and condition as main effects, followed by Tukey's HSD post hoc test for pairwise comparisons between conditions. Conditions sharing a common letter are not significantly different ($P \geq 0.05$); conditions with no shared letters are statistically significant ($P < 0.05$).
