## Supplementary Notes for "Human knee osteoarthritis patient-specific cartilage-on-a-chip model captures donor differences to stressors and treatments"

**Supplementary Note 1. PD-CartChip design validation procedure**

The PD-CartChip platform was developed using a sequential engineering design and validation procedure that combined experimental characterization, analytical modeling, numerical simulation, and mechanical verification (Supplementary Fig. 1). The procedure began with mechanical characterization of native osteoarthritic cartilage explants to establish geometric constraints and compressive loading requirements for device operation. Measurements of explant dimensions defined the required flexure travel range, while unconfined compression testing established the target reaction forces required for the internal flexure arrays.

These experimentally derived design requirements were then used to guide flexure optimization through a multi-stage theoretical screening process. Candidate beam geometries were evaluated using Tresca-based maximum deflection criteria to eliminate designs that could not achieve required travel distances. Remaining candidates were assessed first using linear guided-beam theory to estimate stiffness and force-displacement behavior, and then by bending stress, safety factor, and parasitic shortening calculations to ensure structural integrity. Selected beam configurations were subsequently evaluated using the nonlinear Bi-Beam Constraint Model (Bi-BCM) to quantify geometric stiffening effects and to validate the linear analytical approximations across the intended operating range.

Optimized flexure geometries, identified through analytical screening, were fabricated and incorporated into the PD-CartChip platform. Mechanical performance was experimentally validated using Instron-based force-displacement testing to quantify system stiffness, peak loading capacity, and repeatability. Lastly, optical motion tracking of the carriage and compression paddles during cyclic loading confirmed synchronized motion and verified that the fabricated device reproduced the intended loading mechanics under physiological compression conditions.

Overall, this procedure established a progression from biological characterization, through theoretical and computational optimization, to experimental validation, providing a systematic framework for the design and verification of the PD-CartChip platform.

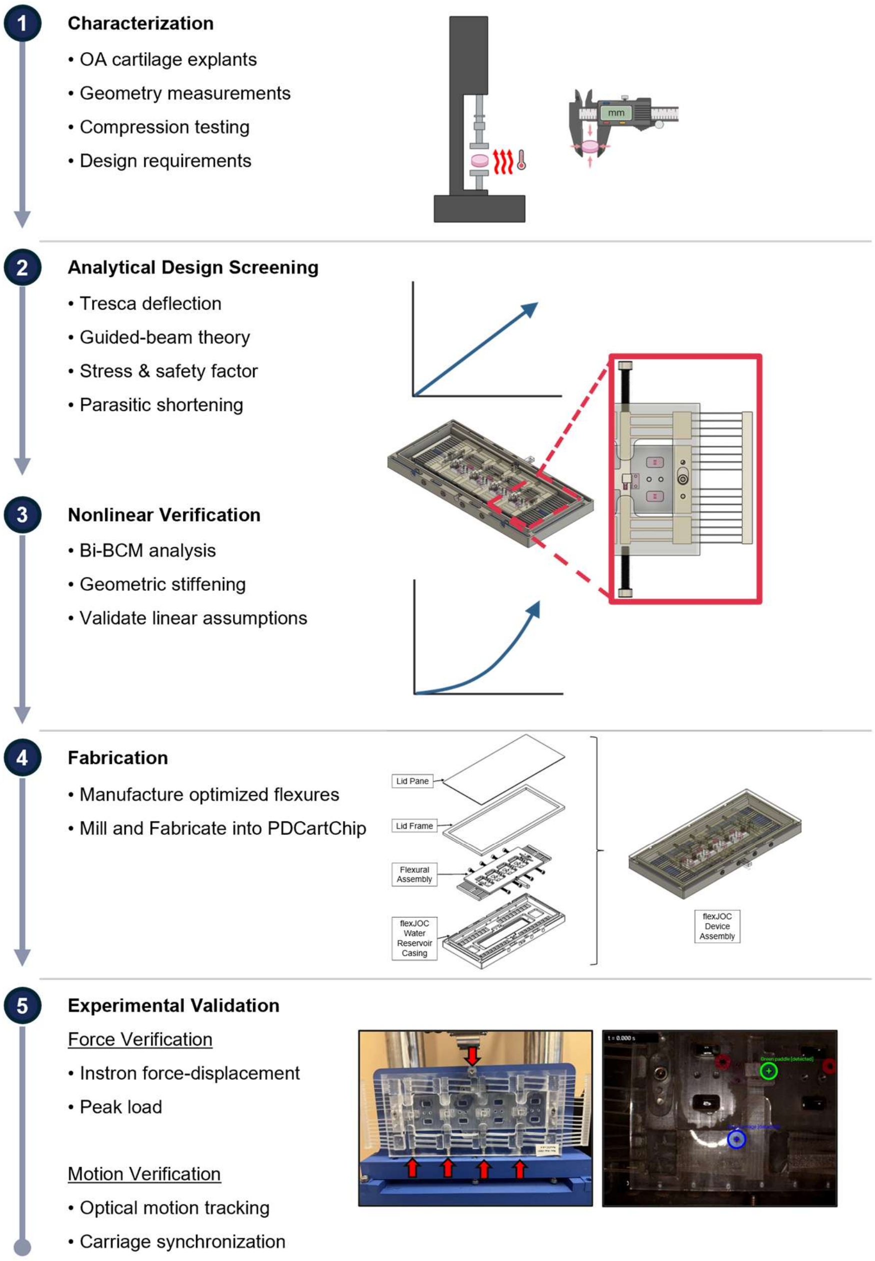

**Supplementary Fig. 1 | PD-CartChip: Overview of design and fabrication procedure**. Step 1 - Characterization: Characterization of cartilage explant peak load response and dimensional variation across donors. Step 2 - Analytical Design Screening: The stepwise screening ideal case analysis completed to determine the final design. Step 3 - Non-Linear Verification: Use of Bi-Beam Constraint model (Bi-BCM) to assess whether non-linear behaviour was a fair assumption for the given translation range. Step 4 - Fabrication: Device fabrication steps. Step 5 - Experimental Validation: Force verification of assembled devices on Instron mechanical tester and motion verification of paddle relative to the carriage translation using a digital microscope camera at 50 fps. Created with Biorender.com.

**Supplementary Note 2. Cartilage explant preparation and mechanical characterization**

**Methods**

Native human osteoarthritic cartilage explants were mechanically characterized to establish the design requirements for the internal flexure arrays (Supplementary Fig. 2). Cylindrical explants (2 mm diameter) were harvested from total knee arthroplasty specimens and equilibrated for 2–3 days before testing. Explant diameter and length were measured using a Keyence VHX-7000 digital microscope, and samples between 1 and 3 mm in length were included for analysis. Stress calculations were normalized using the average explant cross-sectional area. Dimensional variability across donor samples was assessed using one-way ANOVA followed by Tukey's post hoc test when appropriate ($p < 0.05$).

Mechanical characterization was performed using a TA Instruments Q800 Dynamic Mechanical Analyzer under unconfined compression. Samples were pre-equilibrated for 5 min at 37°C in DPBS and compressed to 30% strain at 0.01 mm /s using a 10 mm stainless steel platen following a 0.02 N preload. Force-displacement data were exported using TA Analysis software for subsequent analysis in MATLAB.

Following completion of the initial device optimization, additional testing was performed using a CellScale Univert mechanical tester operating at deformation rates representative of PD-CartChip cyclic loading conditions to confirm that the selected flexure geometry remained appropriate under physiological loading rates using a single donor (N=1), and three explant replicates (n=3).

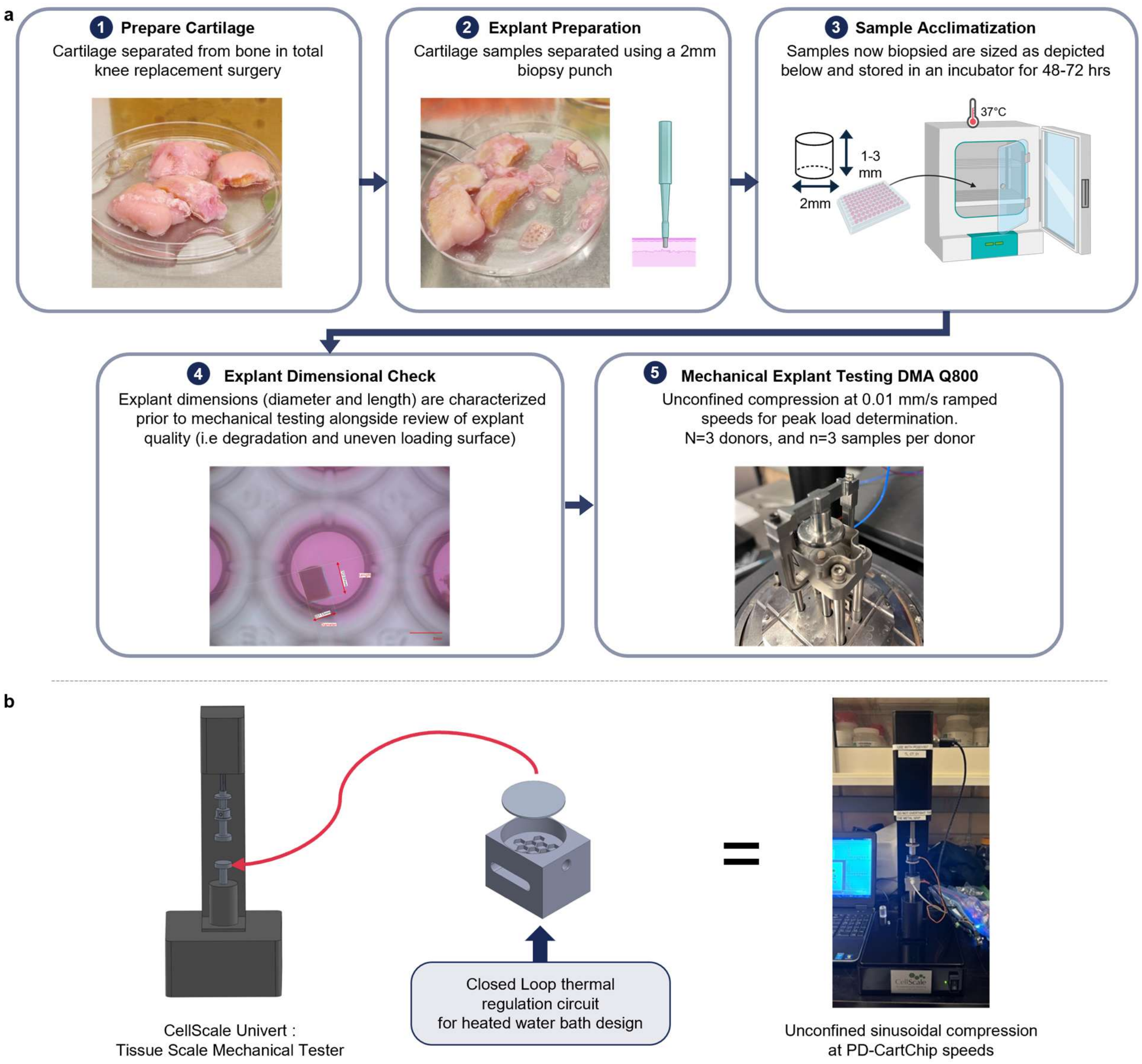

**Supplementary Fig. 2 | Cartilage characterization setups. a,** Primary cartilage characterization workflow illustrating sample preparation, dimensional characterization using the Keyence VHX-7000 digital microscope, and compressive mechanical testing using the TA Instruments Dynamic Mechanical Analyzer (DMA) Q800. Created with Biorender.com. **b,** Additional high deformation rate heated mechanical testing setup: a CellScale Univert Tester coupled with a custom aluminum media bath used for cyclic sinusoidal testing.

**Results**

Under unconfined ramp compression to 30% strain (0.01 mm/s) at 37°C, cartilage explants exhibited characteristic nonlinear increase in compressive force with increasing strain (Supplementary Fig. 3a). Across donors, the average peak load was 3.30 ± 1.09 N, with individual donor means ranging from 2.71 ± 0.96 N to 4.27 ± 0.93 N. Based on these measurements, the expected operational loading range for the PD-CartChip platform was established as approximately 2.2 to 4.3 N. 4.3 N was used as the primary loading target for flexure stiffness.

To assess the relevance of the characterization data to PD-CartChip operation, an additional mechanical test was performed at deformation rates and frequencies comparable to those generated by the device during cyclic loading (i.e., 1 Hz, 0.5 mm/s to 0.9 mm/s). Under sinusoidal compression, cartilage explants exhibited moderately higher peak reaction forces than those measured during the quasi-static ramp tests at 0.01 mm/s during initial cycles, consistent with the viscoelastic rate-dependent behavior of native cartilage (Supplementary Fig. 3b). However, the increase in peak load remained within the operational capacity of the optimized internal flexure arrays and did not require modification of the selected beam geometry. Instead, the slightly greater tissue resistance resulted in a small increase in carriage translation required to achieve the prescribed compressive strain, while maintaining stable device operation. Additionally, after the initial cycle's samples maintain a low operational peak load threshold of 0.56 N for the single donor tested. These base findings indicate that the quasi-static characterization provided a useful baseline for flexure design and that the final PD-CartChip configuration remained suitable for physiological cyclic loading conditions.

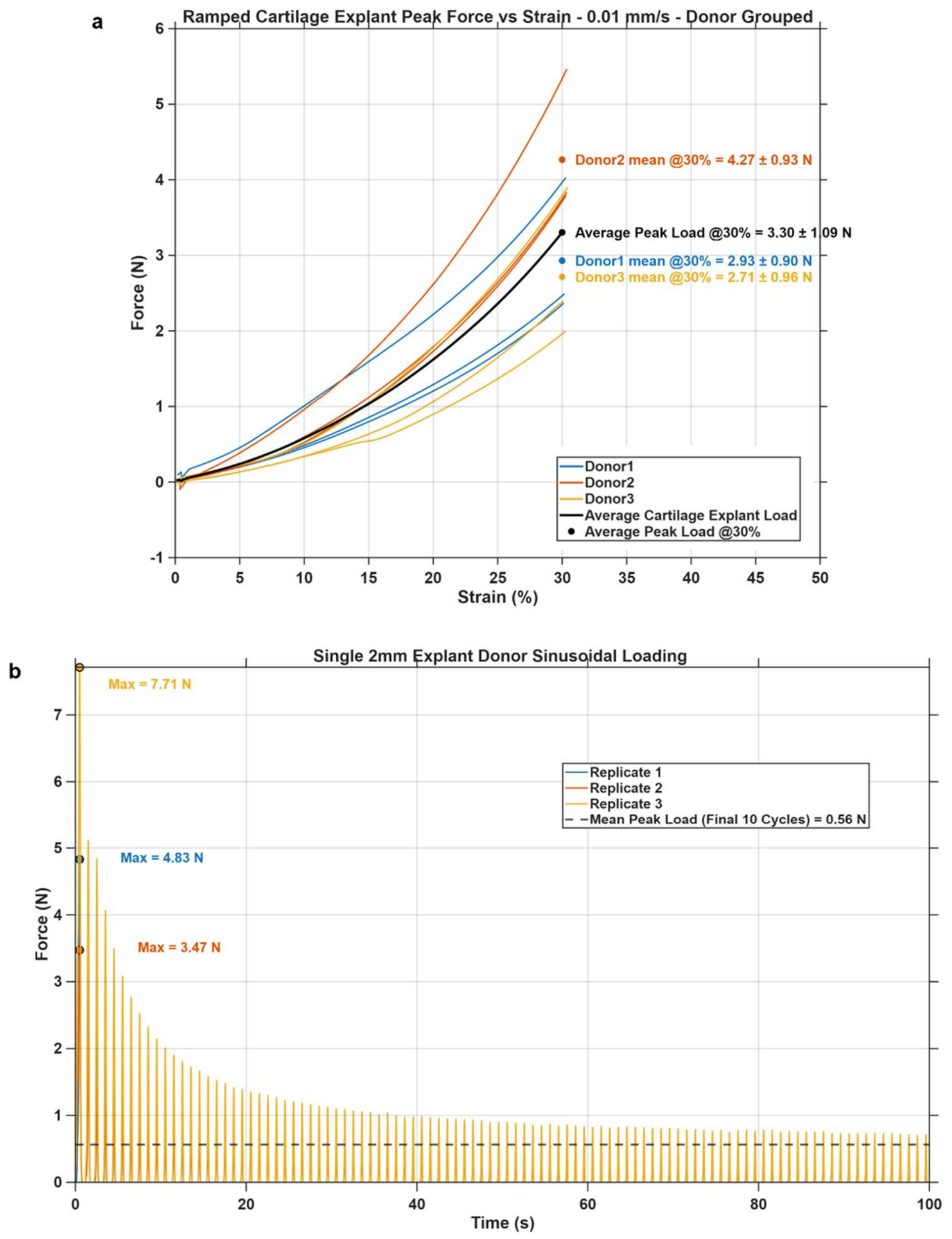

**Supplementary Fig. 3 | Cartilage dimensional and mechanical characterization workflow results. a,** Initial cartilage characterization Dynamic Mechanical Analyzer (DMA) workflow results. Average load–displacement response of 2 mm cartilage explants compressed at a rate of 0.01 mm/s. Solid lines represent the mean response for each donor, with the average peak load highlighted. Three explants were tested per donor (n=3), with three independent donors analyzed (N=3). **b,** Sinusoidal loading Data at 1 Hz tested for one donor (N=1) with three independent replicate explants tested (n=3). Donor 1 to 3 are not the same individuals as Donors 1 to 3 in the main manuscript and annotated in Supplementary Table 1. Tests were carried out on the UniVert Mechanical Tester to capture peak load results on 2 mm explants for 100 cycles.

### Supplementary Note 3. PD-CartChip device design and fabrication

#### Methods

The overall PD-CartChip architecture and fabrication workflow are described in the Main Methods. Briefly, the PD-CartChip platform was designed to deliver cyclic unconfined compressive loading to multiple cartilage explants simultaneously using a single servo-driven actuator. Motion was transmitted through stacked external double-parallelogram flexures to a translating carriage supporting independent internal flexures coupled to acrylic compression paddles. Adjustable limit screws permitted independent strain targeting for explants of varying lengths while maintaining synchronized carriage motion.

The actuator consisted of a high-load servo coupled to a rack-and-pinion mechanism capable of generating 3.5 mm carriage translation. The device comprised of a poly(methyl methacrylate) (PMMA) microwell plate enclosed within an acrylic housing mounted on an acrylic baseplate. Internal and external flexural assemblies were machined from PMMA and integrated with the actuator to provide programmable cyclic loading under explant culture conditions.

#### Results

PD-CartChip Design

The optimized PD-CartChip platform incorporated redesigned internal and external flexure arrays based on experimentally measured cartilage loading requirements. Internal flexures consisted of triple-beam arrays (25 × 5 × 1.5 mm per beam), while external carriage supports comprised sixteen 30 × 5 × 0.5 mm beams arranged in double parallelogram flexure arrays. The redesigned microwell plate accommodated four dynamically loaded explants and two static control wells per loading replicate, with enlarged well spacing and lateral side channels to maintain near-unconfined compression. The completed device integrated a servo-driven rack-and-pinion actuator capable of producing 3.5 mm carriage translation at a maximum loading speed of 7 mm/s.

Flexure Optimization

Successive analytical screening reduced the candidate beam geometries to a limited number of configurations satisfying both displacement and safety requirements. Tresca-based deflection analysis (Supplementary Fig. 4a) identified 25 × 5 × 1.0 mm and 25 × 5 × 1.5 mm internal beams as suitable candidates for the required 1.8 mm travel, while 30 × 5 × 0.5 mm beams satisfied the 3.5 mm travel requirement for the external carriage flexures.

Guided-beam stiffness analysis (Supplementary Fig. 4b) demonstrated an approximately linear increase in stiffness with increasing beam number. Triple 25 × 5 × 1.5 mm beam arrays provided the best compromise between stiffness and allowable deflection and were selected for fabrication.

Maximum predicted bending stresses remained below the PMMA yield strength, corresponding to safety factors of 1.68 for the internal flexures and 2.43 for the external flexures. Estimated parasitic shortening was negligible (~0.065 mm). The final selected internal flexure design was found using this procedure.

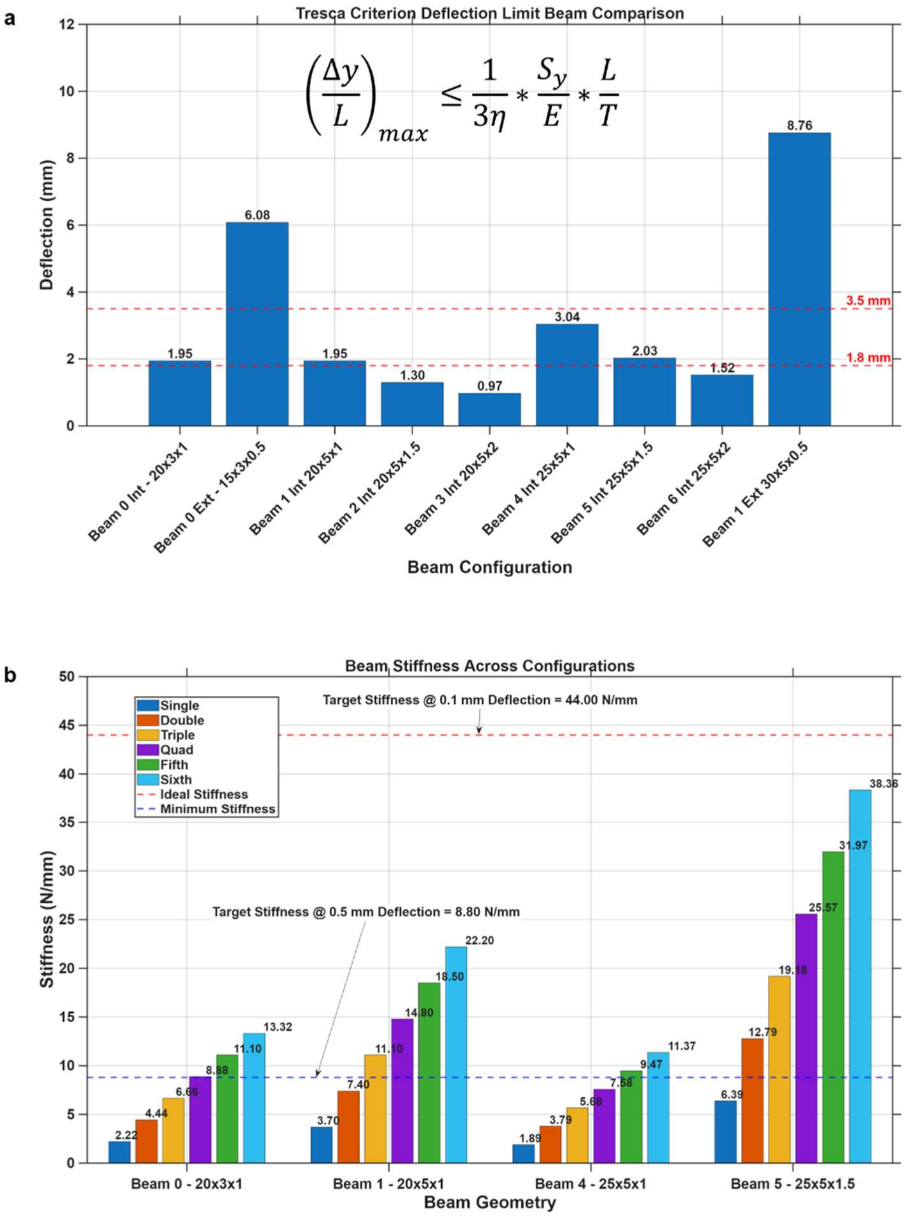

**Supplementary Fig. 4 | Analytical design screening results for flexure selection. a,** Tresca maximum deflection capability comparison across initial beam configurations. Beam alternative number and dimensions listed in the x-axis as "Beam # – Int or Ext – LxHxT". Identifiers are: "Int" = Internal Flexure Option; "Ext" = External flexure option; "LxHxT" = Length x Height x Thickness of beam. Minimum translational distance required by external flexures and internal flexures in the PD-CartChip design is 3.5 mm and 1.8 mm, respectively. **b,** Stiffness evaluation for stacked beam arrangements. Linear stiffness was calculated and assessed for all flexure dimensions surviving the initial deflection screening step and projected based on potential stacked configurations, ranging from 1 to 6 beams on either side of the paddle (2 to 12 beams total). Lower blue dotted line: The minimum target stiffness required for the internal flexure array to resist cartilage peak loads of 4.3 N with a deflection of 0.5 mm. Upper red dotted line: The maximum target stiffness required for the internal flexure array to resist cartilage peak loads of 4.3 N with a deflection of 0.1 mm. In both cases, the carriage would travel this extra distance to allow for the cartilage to hit compression targets; however, the ideal case would be as small as possible.

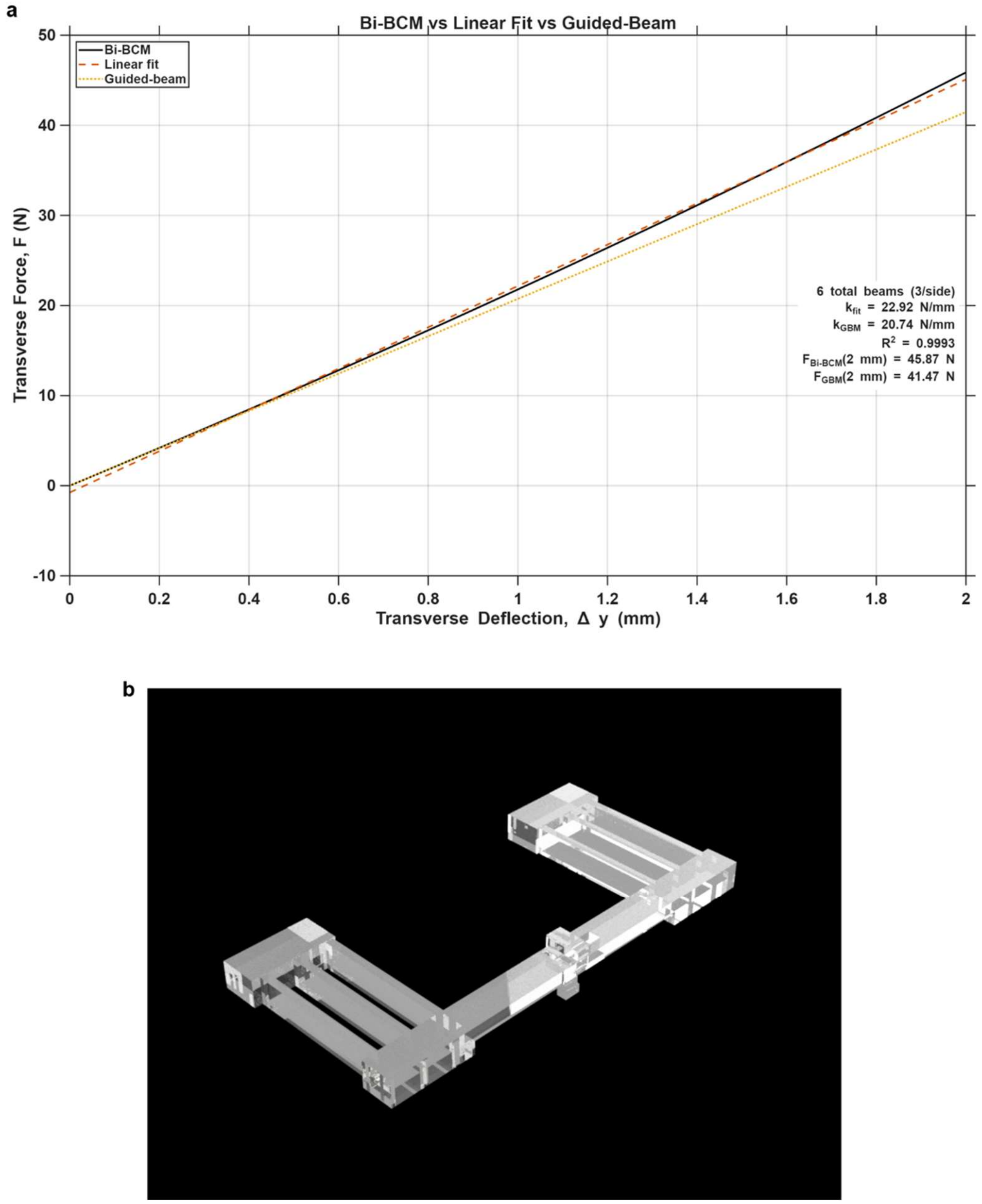

**Supplementary Fig. 5 | Analytical design screening results for flexure selection**. **a,** Non-Linear vs. Linear Comparison of Triple 25 x 5 x 1.5 mm Parallelogram Internal Flexure Configurations. Comparison of the linear regression of the Bi-Beam Constraint Model (Bi-BCM model) and guided-beam model (GBM) predictions for the triple 25 × 5 × 1.5 mm internal flexure configuration (six beams total) over a 0–2 mm deflection range. **b,** Final Selected Internal Flexure Design. Triple stacked 25 x 5 x 1.5 mm flexures on either side of the paddle. Selected as the best configuration based on the balancing of flexible travel range, and resistive stiffness for the 4.3 N peak cartilage load limit collected during characterization.

Comparison of guided-beam theory with the nonlinear Bi-BCM model demonstrated excellent agreement over the operating range (Supplementary Fig. 5a). The Bi-BCM model predicted an effective stiffness of 22.9 N/mm compared with 20.7 N/mm from the guided-beam model, representing approximately a 10% difference. Root mean square error between the analytical models was approximately 10% across the full operating range, confirming that the guided-beam approximation provided an accurate screening method for flexure design.

### Supplementary Note 4. Device validation

#### Methods

Mechanical validation of the completed flexure assemblies was performed using an Instron 5848 Universal Testing system equipped with a 10 kN load cell. Custom 3D-printed fixtures replicated the constraints of the assembled device while force-displacement testing quantified flexure stiffness, linearity, repeatability, and peak loading capacity under both carriage translation and independent internal flexure loading conditions.

To verify synchronized device operation, carriage and paddle motion were quantified using optical motion tracking. Fiduciary markers attached to the carriage and compression paddles were imaged using a Keyence VHX-7000 microscope at 50 fps under brightfield illumination. Marker positions were tracked using a custom Python image-processing pipeline based on HSV color segmentation. The blue marker diameter (5.9 mm) was used to convert pixel measurements to millimetres. Videos longer than 10 seconds were automatically limited to the first 10 seconds, and loading cycles were identified automatically. Hold periods, during which the paddle remained stationary at its mechanical hard stop, were excluded from the synchronization analysis. Paddle synchronization was assessed using a pre-hold trajectory residual, which compares the measured paddle motion with the expected paddle motion based on carriage displacement before the hard stop. Temporal lag, trajectory residuals, peak carriage displacement, and lateral paddle drift were calculated for each loading cycle, averaged within each replicate, and reported across four explant replicates as mean ± standard deviation.

#### Results

Mechanical characterization of whole flexural assemblies

Mechanical testing of fabricated flexural assemblies using an Instron Universal Testing system demonstrated consistent external flexure performance (Supplementary Fig. 6). Under carriage translation alone, the average peak reaction force was 10.92 ± 0.04 N, with minimal variability between assemblies. This means that to translate the carriage 3.5 mm with no engaged strain targets the actuator must apply 11 N (Supplementary Fig. 6a).

When both external and internal flexures were engaged, the average peak system reaction force increased to 101.54 ± 35.28 N, with a maximum measured load of approximately 137 N (Supplementary Fig. 6b). Both assemblies exhibited similar loading profiles despite increased variability at higher displacements.

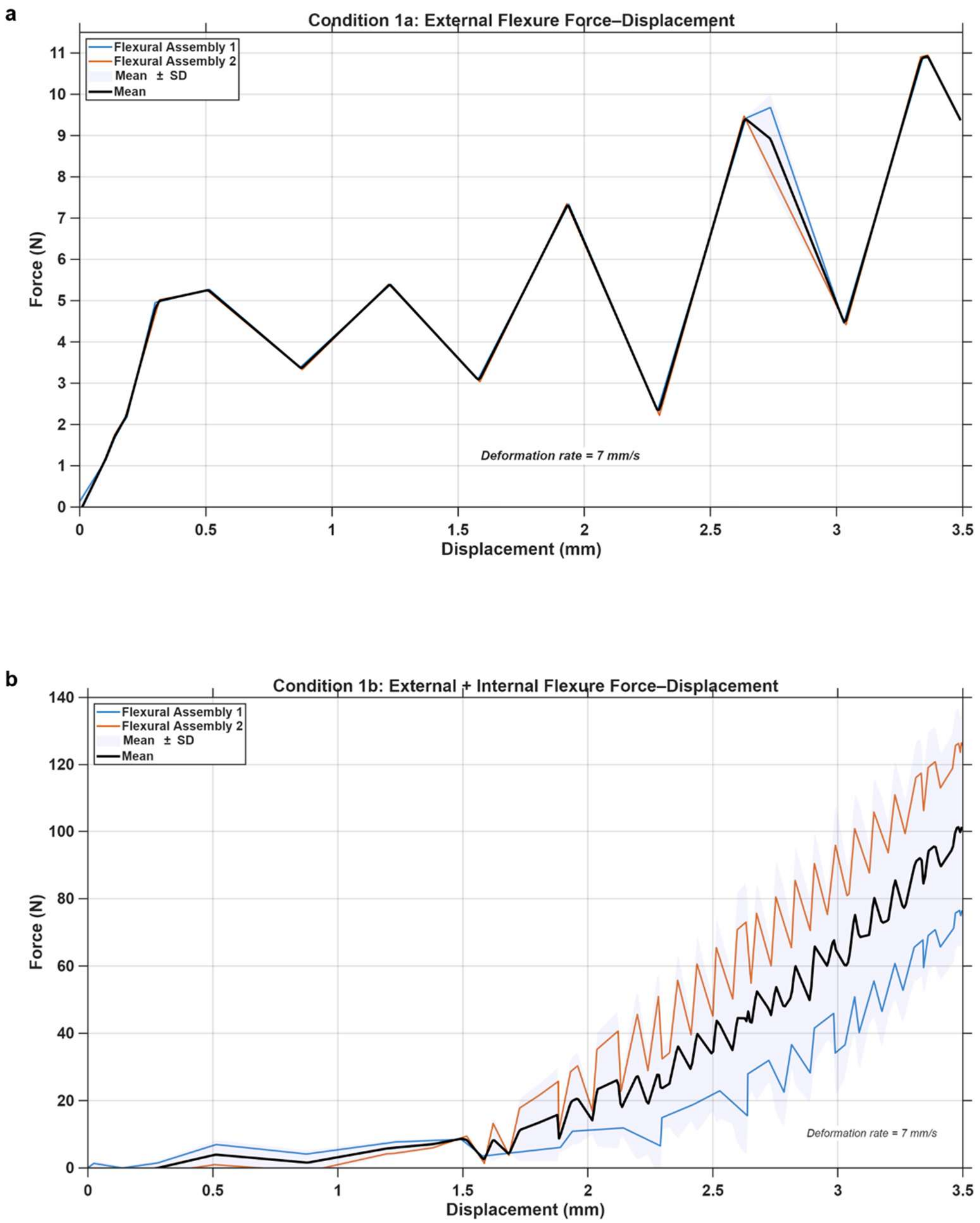

**Supplementary Fig. 6 | Mechanical characterization of the flexural loading assemblies under whole assembly conditions. a,** Force–displacement curves for two external flexure assemblies subjected to a full carriage translation of 3.5 mm at 7mm/s (the estimated maximum speed of the PD-CartChip) (Condition 1a). **b,** Force–displacement curves for two external flexure assemblies subjected to a full carriage translation of 3.5 mm, with all internal flexures simultaneously deflected to a maximum displacement of 2.0 mm (Condition 1b).

Independent mechanical characterization of internal flexure arrays

Independent testing of the internal flexure arrays produced average peak loads of 40.5 N and 55.6 N for the two fabricated assemblies, respectively at a peak translational distance of 2 mm (Supplementary Fig. 7). The experimentally measured average flexure stiffness was 23.8 N/mm, approximately 24% greater than predicted by linear analytical modeling. Assembly 2 exhibited significantly greater stiffness than Assembly 1 (30.72 ± 4.31 vs. 16.77 ± 9.79 N/mm; Student's t-test, p = 0.040).

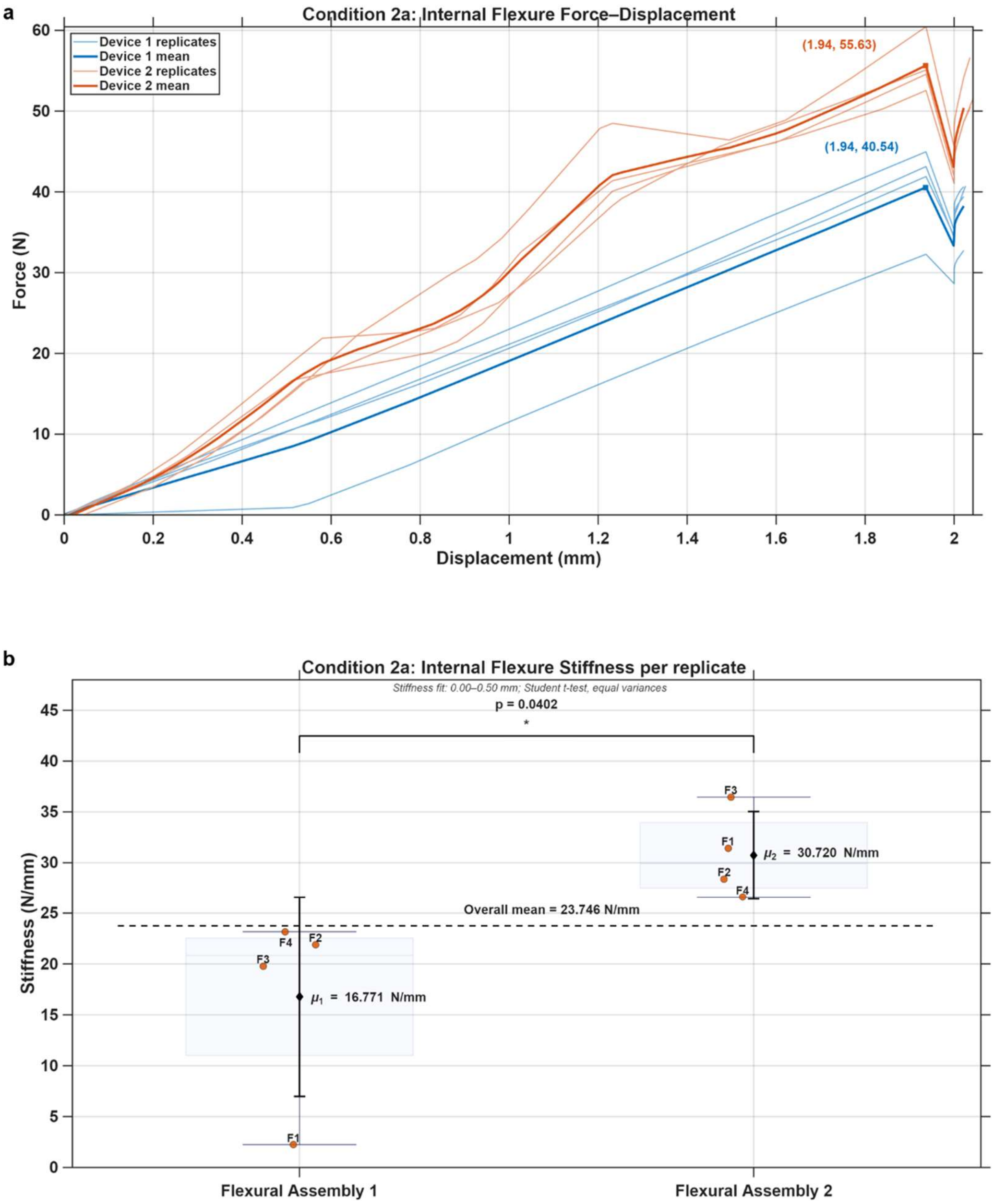

**Supplementary Fig. 7 | Mechanical characterization of the flexural loading assemblies under individual replicate conditions. a,** Force–displacement curves for two internal flexure

assemblies (Condition 2a). Thin lines represent individual replicates (n=4 per device), while bold lines indicate the mean response for each device. Peak force values for the mean curves are annotated at their corresponding displacement. **b,** Internal flexure stiffness calculated from the initial loading region for each replicate. Box plots show the interquartile range with whiskers representing the data spread, orange markers indicate individual replicates, and black diamonds with error bars represent the device mean ± standard deviation. Device 2 exhibited a higher force response and greater stiffness than Device 1 across the loading conditions.

Carriage and paddle motion tracking validation

Motion tracking analysis demonstrated close synchronization between carriage translation and paddle motion during cyclic loading (Supplementary Fig. 8a). Synchronization was assessed by tracking the displacement of the carriage and paddle markers throughout each loading cycle while excluding the hold phase, during which the paddle intentionally remains stationary after contacting its mechanical hard stop screw set to a specified strain target. Rather than comparing the absolute displacement difference between the carriage and paddle, which is expected to increase because the carriage continues translating after the paddle reaches its compression limit, the analysis quantified a pre-hold proportional residual. This residual represents the difference between the measured paddle trajectory and the expected paddle trajectory predicted from the carriage displacement during the active loading phase prior to the hard stop.

Across the four explant replicates, temporal offsets between carriage and paddle motion remained negligible, with no measurable systematic phase shift during active loading (Supplementary Fig. 8b). The mean absolute pre-hold proportional residual ranged from 0.014 to 0.184 mm, with corresponding root-mean-square (RMS) residuals of 0.021 to 0.264 mm, indicating that the paddle closely followed the expected carriage-coupled trajectory throughout compression. The largest instantaneous pre-hold residual observed was 0.572 mm, occurring in the replicate with the greatest compression displacement (3.84 mm), suggesting that increased sample geometry and mechanical compliance contributed to larger transient deviations. Peak lateral paddle drift remained below 120 µm across all four replicates, demonstrating minimal parasitic motion in the horizontal direction. Overall, these results demonstrate that the loading paddle faithfully follows through with the projected carriage motion throughout the active compression phase and that additional carriage translation after the paddle reaches its hard stop reflects continued compression of the sample rather than a loss of synchronization between the loading components.

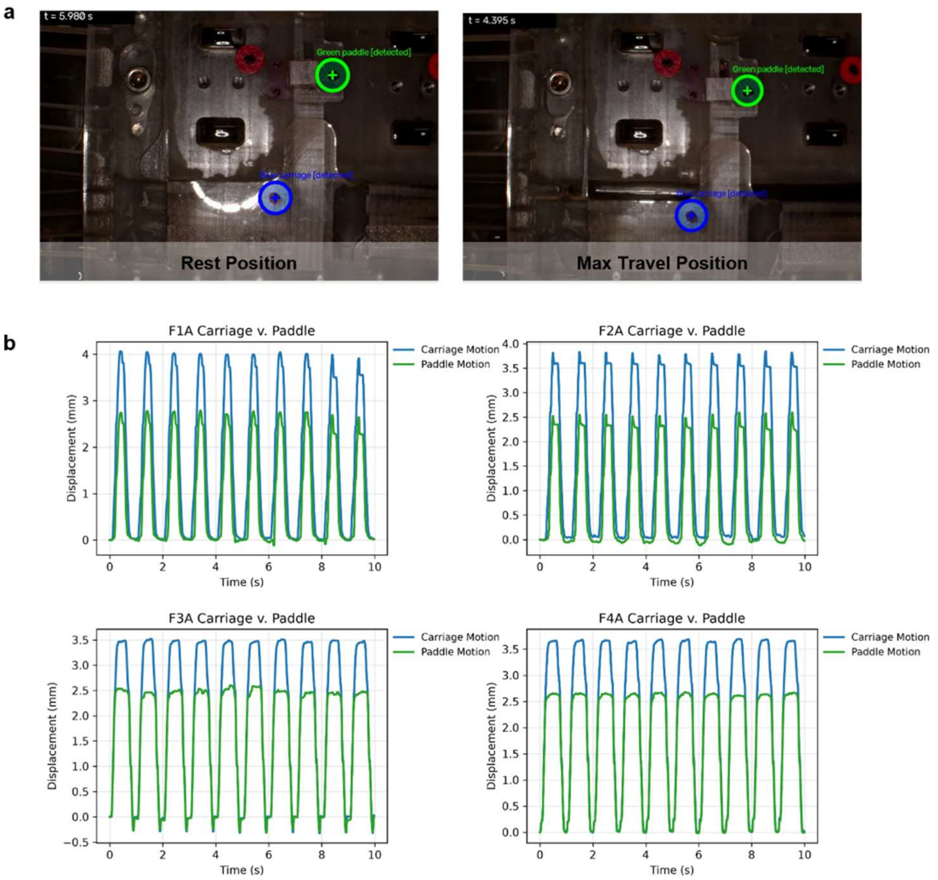

c

| Replicate | Mean absolute pre-hold residual (mm) | RMSE (mm) | Peak residual (mm) | Peak carriage displacement (mm) | Lateral drift (μm) |
|---|---|---|---|---|---|
| F1A | 0.128 | 0.193 | 0.562 | 4.062 | 58.7 |
| F2A | 0.184 | 0.264 | 0.572 | 3.844 | 118.2 |
| F3A | 0.025 | 0.033 | 0.088 | 3.518 | 80.3 |
| F4A | 0.014 | 0.021 | 0.063 | 3.696 | 18.0 |

**Supplementary Fig. 8 | Motion tracking validation of carriage and paddle synchronization during cyclic compression. a,** Representative rest and maximum compression positions showing the blue carriage marker and green paddle marker used for HSV-based motion tracking. Marker outlines indicate the tracked regions used for centroid detection throughout the experiment. **b,** Carriage and paddle displacement versus time for four representative cartilage explant replicates over a 10-second recording. Traces are phase-aligned to facilitate comparison of motion synchronization. Blue traces represent carriage displacement, while green traces represent paddle displacement. The plateau in the paddle trace corresponds to the paddle reaching its mechanical stop, after which continued carriage translation accommodates additional sample compression.
