## Supplementary Tables for "Human knee osteoarthritis patient-specific cartilage-on-a-chip model captures donor differences to stressors and treatments"

Supplementary Table 7. Exploratory correlations: Clinical readouts and experimental responses. 8

**Supplementary Table 1. Patient/Donor characteristics**

| Donor | Symbol | Age (Year) | Sex | BMI (kg/m$^3$) |
|---|---|---|---|---|
| 1 | ● | 72 | Female | 28.8 |
| 2 | ■ | 84 | Male | 29.7 |
| 3 | ▲ | 72 | Female | 37.5 |
| 4 | ◆ | 68 | Male | 28.5 |
| 5 | ◧/□ | 70 | Female | 32.9 |
| 6 | ◭/△ | 62 | Female | 36.3 |
| 7 | ⬣ | 66 | Male | 33.4 |
| 8 | ▼ | 63 | Female | 24.7 |
| 9 | ◓/◯ | 70 | Female | 29.9 |

**Supplementary Table 2. RT-qPCR cartilage forward and reverse primer sequences.**

| Gene | Forward | Reverse |
|---|---|---|
| *B2M* | CTTTCTGGCCTGGAGGCTATC | ACCAGTCCTTGCTGAAAGACAA |
| *ACTB* | CTCACCATGGATGATGATATCGC | AGGAATCCTTCTGACCCATGC |
| *GAPDH* | ACAACTTTGGTATCGTGGAAGG | GCCATCACGCCACAGTTTC |
| *ACAN* | ACTCTGGGTTTTCGTGACTCT | ACACTCAGCGAGTTGTCATGG |
| *SOX9* | ATGCAAGCATGTGTCATCCA | AGGTCTGTCAGTGGGCTGAT |
| *COL1A1* | CCTGGATGCCATCAAAGTCT | CGCCATACTCGAACTGGAAT |
| *COL2A1* | CATGAGGGCGCGGTAGAG | CCAGCCTCCTGGACATCCT |
| *COL10A1* | TGCTGCCACAAATACCCTTT | GTGGACCAGGAGTACCTTGC |
| *PRG4* | AGGCCCCATGTGTTCATGC | GCGCAAAGTAGTCAGTCCATCT |
| *TGFB* | ACTGCGGATCTCTGTGTCAT | AGTAGTGTTCCCCACTGGTCC |
| *TIMP1* | CAATTCCGACCTCGTCATCAG | TATACATCTTGGTCATCTTGATCTCATAAC |
| *ADAMTS4* | CAAGGTCCCATGTGCAACGT | CATCTGCCACCACCAGTGTCT |
| *ADAMTS5* | TGGCTCACGAAATCGGACATT | TGCATTTGGACCAGGGCTTA |
| *MMP1* | CTCAATTTCACTTCTGTTTTCTG | CATCTCTGTCGGCAAATTCGT |
| *MMP3* | CAAAGCTTCAGTGTTGGCTG | GGCCAGGGATTAATGGAGAT |
| *MMP9* | AGACCTGGGCAGATTCCAAAC | CGGCAAGTCTTCCGAGTAGT |
| *MMP13* | TCACCAATTCCTGGGAAGTCT | TCAGGAAACCAGGTCTGGAG |
| *IL6* | TCTCCACAAGCGCCTTCG | CTCAGGGCTGAGATGCCG |
| *IL1B* | CTAAACAGATGAAGTGCTCCT | TAGCTGGATGCCGCCAT |
| *CXCL8* | AAA TTT GGG GTG GAA AGG TT | TCC TGA TTT CTG CAG CTC TGT |
| *PIEZO1* | TCGGACCAGTCTGTGGTCAT | GAGGTAGTCGGCCTCCTCAT |
| *PIEZO2* | GACGGACACAACTTTGAGCCTG | CTGGCTTTGTTGGGCACTCATTG |
| *TRPV4* | CAACGACACCATCCCTGTGC | GGGCAGCTCCCCTCGATA |

**Supplementary Table 3. Statistical analysis across experimental readouts in physiological vs. hyperphysiological compression experiments.**

| Experimental Readout[a] | Time Point | Overall Condition Effect, P[b] | Pairwise Comparison, P [b] | | |
|---|---|---|---|---|---|
| | | | HPC | HPC | PC |
| | | | PC | STATIC | STATIC |
| *ACAN* | Day 14 | 0.012 | 0.636 | 0.011 | 0.079 |
| *SOX9* | Day 14 | 0.380 | 0.456 | 1.000 | 0.441 |
| *COL1A1* | Day 14 | 0.457 | 0.683 | 0.438 | 0.912 |
| *COL2A1* | Day 14 | 0.265 | 0.632 | 0.237 | 0.735 |
| *COL10A1* | Day 14 | 0.016 | 0.316 | 0.232 | 0.012 |
| *PRG4* | Day 14 | <.0001 | 0.272 | <.0001 | <.0001 |
| *TGFB* | Day 14 | 0.136 | 0.200 | 0.998 | 0.178 |
| *TIMP1* | Day 14 | 0.088 | 0.112 | 0.156 | 0.982 |
| *ADAMTS4* | Day 14 | 0.010 | 0.077 | 0.009 | 0.599 |
| *ADAMTS5* | Day 14 | 0.208 | 0.287 | 0.255 | 0.997 |
| *MMP1* | Day 14 | 0.283 | 0.823 | 0.260 | 0.573 |
| *MMP3* | Day 14 | 0.001 | 0.540 | 0.001 | 0.011 |
| *MMP9* | Day 14 | 0.402 | 0.846 | 0.697 | 0.373 |
| *MMP13* | Day 14 | 0.386 | 0.999 | 0.465 | 0.443 |
| *IL6* | Day 14 | 0.061 | 0.901 | 0.152 | 0.067 |
| *IL1B* | Day 14 | 0.955 | 0.954 | 0.997 | 0.975 |
| *CXCL8* | Day 14 | 0.001 | 0.634 | 0.009 | 0.001 |
| *PIEZO1* | Day 14 | 0.001 | 0.775 | 0.006 | 0.001 |
| *PIEZO2* | Day 14 | <.0001 | 0.151 | <.0001 | 0.007 |
| *TRPV4* | Day 14 | 0.039 | 0.093 | 0.050 | 0.947 |
| sGAGs | Day 3 | <.0001 | <.0001 | 0.337 | <.0001 |
| sGAGs | Day 7 | 0.896 | 0.972 | 0.886 | 0.968 |
| sGAGs | Day 10 | 0.001 | 0.016 | 0.672 | 0.002 |
| sGAGs | Day 14 | 0.004 | 0.100 | 0.003 | 0.366 |
| TIMP-1 | Day 7 | 0.018 | 0.740 | 0.088 | 0.017 |
| TIMP-1 | Day 14 | 0.410 | 0.729 | 0.379 | 0.818 |
| SafO | Day 14 | 0.309 | 0.784 | 0.283 | 0.615 |

[a]Experimental readouts included gene expression, sulfated glycosaminoglycans (sGAGs), tissue inhibitor of metalloproteinases 1 (TIMP-1), and Safranin-O proteoglycan staining (SafO).

[b]Statistical significance was defined as $P < 0.05$.

**Supplementary Table 4. Statistical analysis across experimental readouts in inflammatory vs. hyperphysiological compression experiments.**

| Experimental Readout[a] | Time Point | Overall Condition Effect, P[b] | Pairwise Comparison, P [b] | | |
|---|---|---|---|---|---|
| | | | INF HPC | HPC STATIC | INF STATIC |
| *ACAN* | Day 14 | <.0001 | <.0001 | <.0001 | <.0001 |
| *SOX9* | Day 14 | <.0001 | <.0001 | 0.407 | <.0001 |
| *COL1A1* | Day 14 | <.0001 | <.0001 | 0.245 | <.0001 |
| *COL2A1* | Day 14 | <.0001 | <.0001 | 0.002 | <.0001 |
| *COL10A1* | Day 14 | <.0001 | <.0001 | 0.017 | <.0001 |
| *PRG4* | Day 14 | <.0001 | <.0001 | <.0001 | <.0001 |
| *TGFB* | Day 14 | <.0001 | <.0001 | 0.897 | <.0001 |
| *TIMP1* | Day 14 | 0.0104 | 0.022 | 0.024 | 0.999 |
| *ADAMTS4* | Day 14 | <.0001 | 0.006 | 0.001 | <.0001 |
| *ADAMTS5* | Day 14 | <.0001 | <.0001 | <.0001 | <.0001 |
| *MMP1* | Day 14 | <.0001 | <.0001 | 0.173 | <.0001 |
| *MMP3* | Day 14 | <.0001 | <.0001 | <.0001 | <.0001 |
| *MMP9* | Day 14 | 0.0006 | 0.001 | 0.843 | 0.005 |
| *MMP13* | Day 14 | <.0001 | <.0001 | <.0001 | <.0001 |
| *IL6* | Day 14 | <.0001 | <.0001 | 0.908 | <.0001 |
| *IL1B* | Day 14 | <.0001 | <.0001 | 0.986 | <.0001 |
| *IL8* | Day 14 | <.0001 | <.0001 | 0.044 | <.0001 |
| *PIEZO1* | Day 14 | <.0001 | <.0001 | 0.0002 | 0.006 |
| *PIEZO2* | Day 14 | <.0001 | <.0001 | 0.015 | <.0001 |
| *TRPV4* | Day 14 | <.0001 | <.0001 | 0.518 | 0.0002 |
| TIMP-1 | Day 7 | <.0001 | <.0001 | 0.0001 | <.0001 |
| TIMP-1 | Day 14 | 0.0365 | 0.290 | 0.028 | 0.505 |
| Total MMP-13 | Day 7 | <.0001 | <.0001 | 0.134 | <.0001 |
| Total MMP-13 | Day 14 | <.0001 | <.0001 | 0.181 | <.0001 |
| sGAGs | Day 3 | <.0001 | <.0001 | 0.997 | <.0001 |
| sGAGs | Day 7 | <.0001 | <.0001 | 0.275 | <.0001 |
| sGAGs | Day 10 | <.0001 | <.0001 | 0.114 | <.0001 |
| sGAGs | Day 14 | <.0001 | <.0001 | 0.003 | <.0001 |
| SafO | Day 14 | <.0001 | 0.0003 | 0.082 | <.0001 |

[a]Experimental readouts included gene expression, sulfated glycosaminoglycans (sGAGs), tissue inhibitor of metalloproteinases 1 (TIMP-1), total matrix metalloproteinase 13 (Total MMP-13), and Safranin-O proteoglycan staining (SafO).
[b]Statistical significance was defined as $P < 0.05$.

**Supplementary Table 5. Statistical analysis across experimental readouts in dexamethasone treatment experiments.**

| Experimental Readout[a] | Time Point | Overall Condition Effect, P[b] | Pairwise Comparison, P[b] | | | | | | | | | |
|---|---|---|---|---|---|---|---|---|---|---|---|---|
| | | | HPC | HPC | HPC | HPC | HPC+ DEX | HPC+ DEX | HPC+ DEX | INF | INF | INF+ DEX |
| | | | HPC+ DEX | INF | INF+ DEX | STAT-IC | INF | INF+ DEX | STAT-IC | INF+ DEX | STAT-IC | STAT-IC |
| *ACAN* | Day 14 | <.0001 | 0.008 | <.0001 | <.0001 | <.0001 | <.0001 | <.0001 | 0.717 | 0.032 | <.0001 | <.0001 |
| *SOX9* | Day 14 | <.0001 | 0.340 | 0.035 | 0.161 | 0.405 | 0.0001 | 0.001 | 1.000 | 0.963 | 0.0002 | 0.002 |
| *COL1A1* | Day 14 | <.0001 | <.0001 | <.0001 | <.0001 | 0.462 | 0.864 | 0.870 | 0.0003 | 0.309 | 0.007 | <.0001 |
| *COL2A1* | Day 14 | <.0001 | 0.0001 | <.0001 | <.0001 | 0.073 | <.0001 | <.0001 | <.0001 | 0.0004 | <.0001 | <.0001 |
| *COL10A1* | Day 14 | <.0001 | <.0001 | <.0001 | <.0001 | 0.142 | <.0001 | <.0001 | 0.071 | 1.000 | <.0001 | <.0001 |
| *PRG4* | Day 14 | <.0001 | <.0001 | <.0001 | <.0001 | <.0001 | <.0001 | 0.084 | 0.914 | <.0001 | <.0001 | 0.411 |
| *TGFB* | Day 14 | <.0001 | 0.488 | 0.0002 | <.0001 | 1.000 | 0.029 | <.0001 | 0.478 | 0.008 | 0.0002 | <.0001 |
| *TIMP1* | Day 14 | 0.4529 | 0.979 | 0.479 | 0.998 | 0.708 | 0.819 | 0.999 | 0.954 | 0.670 | 0.996 | 0.867 |
| *ADAMTS4* | Day 14 | <.0001 | <.0001 | 0.003 | 0.575 | 0.158 | <.0001 | <.0001 | 0.065 | 0.132 | <.0001 | 0.003 |
| *ADAMTS5* | Day 14 | <.0001 | 0.016 | <.0001 | 0.574 | 0.003 | <.0001 | 0.0002 | 0.978 | <.0001 | <.0001 | <.0001 |
| *MMP1* | Day 14 | <.0001 | <.0001 | <.0001 | <.0001 | 0.638 | <.0001 | <.0001 | <.0001 | 0.001 | <.0001 | <.0001 |
| *MMP3* | Day 14 | <.0001 | 0.986 | <.0001 | 0.003 | 0.019 | <.0001 | 0.012 | 0.004 | 0.0001 | <.0001 | <.0001 |
| *MMP9* | Day 14 | 0.0726 | 1.000 | 0.096 | 0.976 | 1.000 | 0.107 | 0.983 | 1.000 | 0.300 | 0.152 | 0.996 |
| *MMP13* | Day 14 | <.0001 | <.0001 | <.0001 | 0.997 | 0.0002 | <.0001 | <.0001 | <.0001 | <.0001 | <.0001 | <.0001 |
| *IL6* | Day 14 | <.0001 | 0.005 | <.0001 | <.0001 | 0.942 | <.0001 | <.0001 | 0.037 | <.0001 | <.0001 | <.0001 |
| *IL1B* | Day 14 | <.0001 | 0.435 | <.0001 | 0.761 | 0.738 | <.0001 | 0.984 | 0.988 | <.0001 | <.0001 | 1.000 |
| *IL8* | Day 14 | <.0001 | 0.010 | <.0001 | <.0001 | 0.102 | <.0001 | <.0001 | 0.895 | 0.0003 | <.0001 | <.0001 |
| *PIEZO1* | Day 14 | <.0001 | 0.218 | 0.005 | <.0001 | 0.349 | 0.543 | 0.012 | 0.999 | 0.366 | 0.376 | 0.005 |
| *PIEZO2* | Day 14 | <.0001 | 0.0001 | <.0001 | <.0001 | 0.750 | 1.000 | 0.011 | <.0001 | 0.015 | <.0001 | <.0001 |
| *TRPV4* | Day 14 | <.0001 | 1.000 | 0.0004 | 0.002 | 0.997 | 0.0003 | 0.002 | 0.993 | 0.983 | 0.001 | 0.006 |
| TIMP-1 | Day 7 | <.0001 | <.0001 | 0.0003 | <.0001 | 0.002 | 0.867 | 0.977 | <.0001 | 0.995 | <.0001 | <.0001 |
| TIMP-1 | Day 14 | <.0001 | 0.0002 | 0.438 | 0.835 | 0.335 | <.0001 | <.0001 | <.0001 | 0.964 | 1.000 | 0.916 |
| Total | Day 7 | <.0001 | <.0001 | <.0001 | 1.000 | 0.046 | <.0001 | <.0001 | <.0001 | <.0001 | <.0001 | 0.059 |

| | | | | | | | | | | | | |
|---|---|---|---|---|---|---|---|---|---|---|---|---|
| MMP-13 Total MMP-13 | Day 14 | <.0001 | <.0001 | <.0001 | 0.033 | 0.107 | <.0001 | <.0001 | <.0001 | <.0001 | <.0001 | 0.988 |
| sGAGs | Day 3 | <.0001 | 0.935 | <.0001 | <.0001 | 0.956 | <.0001 | <.0001 | 1.000 | 0.001 | <.0001 | <.0001 |
| sGAGs | Day 7 | <.0001 | 0.019 | <.0001 | 0.648 | 0.697 | <.0001 | 0.0001 | 0.369 | <.0001 | <.0001 | 0.065 |
| sGAGs | Day 10 | <.0001 | 0.004 | <.0001 | 0.080 | 0.015 | <.0001 | <.0001 | 0.996 | <.0001 | <.0001 | <.0001 |
| sGAGs | Day 14 | <.0001 | 0.621 | <.0001 | 0.001 | 0.665 | <.0001 | <.0001 | 1.000 | 0.003 | <.0001 | <.0001 |
| SafO | Day 14 | 0.0149 | 1.000 | 0.187 | 0.882 | 0.666 | 0.124 | 0.780 | 0.790 | 0.681 | 0.009 | 0.179 |

[a]Experimental readouts included gene expression, sulfated glycosaminoglycans (sGAGs), tissue inhibitor of metalloproteinases 1 (TIMP-1), total matrix metalloproteinase 13 (Total MMP-13), and Safranin-O proteoglycan staining (SafO).

[b]Statistical significance was defined as $P < 0.05$.

**Supplementary Table 6. Exploratory correlations: Experimental readouts in stressor-specific models.**

| Stressor | Experimental Readout 1[a] | Experimental Readout 2[a] | Spearman's ρ | FDR-adjusted P value[b] |
|---|---|---|---|---|
| HPC | Anabolic GeneEx | sGAG_d14 | 1 | 0.001 |
| HPC | Total_MMP13_d14 | SafO | -1 | 0.001 |
| HPC | Anabolic GeneEx | TIMP1_d14 | 0.8 | 0.480 |
| HPC | TIMP1_d14 | sGAG_d14 | 0.8 | 0.554 |
| HPC | Catabolic GeneEx | Anabolic GeneEx | -0.8 | 0.720 |
| HPC | Catabolic GeneEx | sGAG_d14 | -0.8 | 0.800 |
| HPC | Catabolic GeneEx | TIMP1_d14 | -0.6 | 0.800 |
| HPC | TIMP1_d14 | SafO | 0.2 | 0.800 |
| HPC | Catabolic GeneEx | SafO | 0.2 | 0.800 |
| HPC | Catabolic GeneEx | Total_MMP13_d14 | -0.2 | 0.873 |
| HPC | Anabolic GeneEx | Total_MMP13_d14 | 0.4 | 0.900 |
| HPC | Total_MMP13_d14 | TIMP1_d14 | -0.2 | 0.929 |
| HPC | Total_MMP13_d14 | sGAG_d14 | 0.4 | 0.939 |
| HPC | sGAG_d14 | SafO | -0.4 | 1.000 |
| HPC | Anabolic GeneEx | SafO | -0.4 | 1.000 |
| INF | Total_MMP13_d14 | SafO | -1 | 0.004 |
| INF | Anabolic GeneEx | Total_MMP13_d14 | -0.8 | 0.720 |
| INF | Catabolic GeneEx | Anabolic GeneEx | -0.8 | 0.720 |
| INF | Catabolic GeneEx | Total_MMP13_d14 | 0.4 | 0.800 |
| INF | Catabolic GeneEx | sGAG_d14 | 0.4 | 0.831 |
| INF | Catabolic GeneEx | SafO | -0.4 | 0.864 |
| INF | Anabolic GeneEx | SafO | 0.8 | 0.900 |
| INF | Anabolic GeneEx | sGAG_d14 | -0.2 | 0.929 |
| INF | Total_MMP13_d14 | TIMP1_d14 | -0.4 | 0.982 |
| INF | sGAG_d14 | SafO | -0.4 | 1.000 |
| INF | Anabolic GeneEx | TIMP1_d14 | 0 | 1.000 |
| INF | TIMP1_d14 | SafO | 0.4 | 1.000 |
| INF | TIMP1_d14 | sGAG_d14 | 0.4 | 1.000 |
| INF | Total_MMP13_d14 | sGAG_d14 | 0.4 | 1.000 |
| INF | Catabolic GeneEx | TIMP1_d14 | 0.6 | 1.000 |

[a]Experimental readouts included anabolic and catabolic pathway-related gene expression (GeneEx), sulfated glycosaminoglycans (sGAGs), tissue inhibitor of metalloproteinases 1 (TIMP1), total matrix metalloproteinase 13 (Total_MMP13), and Safranin-O proteoglycan staining (SafO) at day 14 (d14).

[b]P values were adjusted for multiple comparisons using the Benjamini–Hochberg false discovery rate (FDR) procedure. Statistical significance was defined as FDR-adjusted $P < 0.05$.

**Supplementary Table 7. Exploratory correlations: Clinical readouts and experimental responses.**

| Stressor+ Treatment | Clinical Readout[a] | Experimental Readout[b] | Spearman's ρ | FDR-adjusted P value[c] |
|---|---|---|---|---|
| HPC+DEX | Pain | Anabolic GeneEx | -1.00 | 0.003 |
| HPC+DEX | Med_NSAID_Freq | Catabolic GeneEx | -0.77 | 0.696 |
| HPC+DEX | QOL | Total_MMP13_d14 | -0.80 | 0.700 |
| HPC+DEX | Med_NSAID_Freq | Total_MMP13_d14 | -0.77 | 0.717 |
| HPC+DEX | Med_NSAID_Freq | SafO | 0.77 | 0.740 |
| HPC+DEX | Symptom | Anabolic GeneEx | 0.74 | 0.744 |
| HPC+DEX | Sport | TIMP1_d14 | 0.80 | 0.750 |
| HPC+DEX | Symptom | TIMP1_d14 | 0.95 | 0.770 |
| HPC+DEX | ADL | Anabolic GeneEx | -0.80 | 0.840 |
| HPC+DEX | QOL | Catabolic GeneEx | -0.40 | 0.875 |
| HPC+DEX | QOL | TIMP1_d14 | 0.40 | 0.875 |
| HPC+DEX | ADL | Total_MMP13_d14 | -0.80 | 0.875 |
| HPC+DEX | QOL | sGAG_d14 | -0.20 | 0.875 |
| HPC+DEX | Sport | Catabolic GeneEx | -0.20 | 0.875 |
| HPC+DEX | Sport | SafO | 0.20 | 0.884 |
| HPC+DEX | ADL | sGAG_d14 | 0.20 | 0.894 |
| HPC+DEX | QOL | SafO | 0.40 | 0.900 |
| HPC+DEX | Pain | Catabolic GeneEx | -0.20 | 0.903 |
| HPC+DEX | Sport | Total_MMP13_d14 | -0.40 | 0.913 |
| HPC+DEX | Pain | SafO | 0.20 | 0.913 |
| HPC+DEX | Sport | sGAG_d14 | -0.40 | 0.926 |
| HPC+DEX | Symptom | Total_MMP13_d14 | -0.11 | 0.949 |
| HPC+DEX | Symptom | Catabolic GeneEx | 0.11 | 0.949 |
| HPC+DEX | ADL | Catabolic GeneEx | -0.40 | 0.955 |
| HPC+DEX | Symptom | SafO | -0.11 | 0.959 |
| HPC+DEX | ADL | TIMP1_d14 | -0.40 | 0.969 |
| HPC+DEX | Med_NSAID_Freq | Anabolic GeneEx | 0.26 | 0.974 |
| HPC+DEX | Symptom | sGAG_d14 | -0.63 | 0.989 |
| HPC+DEX | Med_NSAID_Freq | TIMP1_d14 | 0.26 | 0.999 |
| HPC+DEX | Pain | TIMP1_d14 | -0.80 | 1.000 |
| HPC+DEX | Pain | Total_MMP13_d14 | -0.40 | 1.000 |
| HPC+DEX | QOL | Anabolic GeneEx | 0.00 | 1.000 |
| HPC+DEX | Med_NSAID_Freq | sGAG_d14 | 0.26 | 1.000 |
| HPC+DEX | ADL | SafO | 0.40 | 1.000 |
| HPC+DEX | Pain | sGAG_d14 | 0.40 | 1.000 |
| HPC+DEX | Sport | Anabolic GeneEx | 0.60 | 1.000 |
| INF+DEX | ADL | Anabolic GeneEx | -1.00 | 0.004 |
| INF+DEX | Med_NSAID_Freq | Total_MMP13_d14 | -0.77 | 0.789 |
| INF+DEX | Symptom | Total_MMP13_d14 | -0.74 | 0.834 |

| | | | | |
|---|---|---|---|---|
| INF+DEX | QOL | Catabolic GeneEx | 0.80 | 0.840 |
| INF+DEX | Med_NSAID_Freq | TIMP1_d14 | 0.77 | 0.845 |
| INF+DEX | QOL | TIMP1_d14 | 0.80 | 0.875 |
| INF+DEX | QOL | Total_MMP13_d14 | -0.40 | 0.913 |
| INF+DEX | QOL | sGAG_d14 | 0.80 | 0.913 |
| INF+DEX | Sport | SafO | 0.20 | 0.923 |
| INF+DEX | QOL | SafO | 0.40 | 0.940 |
| INF+DEX | Sport | Catabolic GeneEx | 0.40 | 0.955 |
| INF+DEX | Sport | Total_MMP13_d14 | -0.80 | 0.955 |
| INF+DEX | Pain | SafO | 0.20 | 0.966 |
| INF+DEX | Sport | TIMP1_d14 | 0.40 | 0.969 |
| INF+DEX | Symptom | TIMP1_d14 | 0.11 | 0.978 |
| INF+DEX | Sport | sGAG_d14 | 0.40 | 0.984 |
| INF+DEX | Symptom | sGAG_d14 | 0.32 | 0.997 |
| INF+DEX | Symptom | Catabolic GeneEx | 0.32 | 0.997 |
| INF+DEX | Med_NSAID_Freq | Catabolic GeneEx | 0.26 | 0.999 |
| INF+DEX | Pain | Anabolic GeneEx | -0.80 | 1.000 |
| INF+DEX | QOL | Anabolic GeneEx | -0.60 | 1.000 |
| INF+DEX | Med_NSAID_Freq | SafO | -0.26 | 1.000 |
| INF+DEX | Med_NSAID_Freq | Anabolic GeneEx | -0.26 | 1.000 |
| INF+DEX | Sport | Anabolic GeneEx | 0.00 | 1.000 |
| INF+DEX | Symptom | Anabolic GeneEx | 0.21 | 1.000 |
| INF+DEX | Med_NSAID_Freq | sGAG_d14 | 0.26 | 1.000 |
| INF+DEX | Symptom | SafO | 0.32 | 1.000 |
| INF+DEX | ADL | SafO | 0.40 | 1.000 |
| INF+DEX | ADL | Total_MMP13_d14 | 0.40 | 1.000 |
| INF+DEX | Pain | sGAG_d14 | 0.40 | 1.000 |
| INF+DEX | Pain | TIMP1_d14 | 0.40 | 1.000 |
| INF+DEX | Pain | Catabolic GeneEx | 0.40 | 1.000 |
| INF+DEX | ADL | sGAG_d14 | 0.80 | 1.000 |
| INF+DEX | ADL | TIMP1_d14 | 0.80 | 1.000 |
| INF+DEX | ADL | Catabolic GeneEx | 0.80 | 1.000 |
| INF+DEX | Pain | Total_MMP13_d14 | 0.80 | 1.000 |

[a]Clinical readouts comprised non-steroidal anti-inflammatory drugs usage frequency (Med_NSAID_Freq) and Knee Injury and Osteoarthritis Outcome Score (KOOS) subscales, including pain, symptoms, function in daily living (ADL), function in sport and recreation (Sport), and knee-related quality of life (QOL). KOOS sub scores were inverted such that higher scores indicate greater severity of knee problems.

[b]Experimental readouts included anabolic and catabolic pathway-related gene expression (GeneEx), sulfated glycosaminoglycans (sGAGs), tissue inhibitor of metalloproteinases 1 (TIMP1), total matrix metalloproteinase 13 (Total_MMP13), and Safranin-O proteoglycan staining (SafO) at day 14 (d14).

[c]P values were adjusted for multiple comparisons using the Benjamini–Hochberg false discovery rate (FDR) procedure. Statistical significance was defined as FDR-adjusted $P < 0.05$.
